# Targeting Liver Epsins Ameliorates Dyslipidemia in Atherosclerosis through Inhibition of Proprotein Convertase Subtilisin/Kexin Type 9-Mediated Low-density Lipoprotein Receptor Degradation

**DOI:** 10.1101/2024.08.26.609742

**Authors:** Bo Zhu, Krishan Gupta, Kui Cui, Beibei Wang, Marina V. Malovichko, Kathryn Li, Hao Wu, Kulandai Samy Arulsamy, Bandana Singh, Jianing Gao, Scott Wong, Patricia G Yancey, Douglas B. Cowan, Da-Zhi Wang, Sudha Biddinger, MacRae F. Linton, Wenyi Wei, Sanjay Srivastava, Chris A. Bashur, Changcheng Zhou, Kaifu Chen, Hong Chen

## Abstract

**Background:** The low-density lipoprotein receptor (LDLR) in the liver plays a crucial role in clearing low-density lipoprotein cholesterol (LDL-C) from the bloodstream. This process takes place mainly in the liver. Under atherogenic conditions, Proprotein Convertase Subtilisin/Kexin Type 9 (PCSK9), secreted by the liver, binds to LDLR on hepatocytes, preventing its recycling and enhancing its lysosomal degradation. This process reduces LDL-C clearance, promoting hypercholesterolemia. Epsins, a family of ubiquitin-binding endocytic adaptors, are key regulators of atherogenesis in lesional cells, including endothelial cells and macrophages. Given epsins’ canonical role in regulating endocytosis of cell surface receptors, we aimed to determine whether and how liver epsins contribute to PCSK9-mediated LDLR endocytosis and degradation, thereby impairing LDL-C clearance and accelerating atherosclerosis.

**Methods:** Liver-specific epsin knockout (Liver-DKO) atherosclerotic models were generated in ApoE^-/-^ and PCSK9-AAV8-induced atheroprone mice fed on a Western diet. We utilized single-cell RNA sequencing, along with molecular, cellular, and biochemical analyses, to investigate the physiological role of liver epsins in PCSK9-mediated LDLR degradation. Additionally, we explored the therapeutic potential of nanoparticle-encapsulated siRNAs targeting epsins 1 and 2 in ApoE^-/-^ mice with established atherosclerosis.

**Results:** Western diet (WD)-induced atherosclerosis was significantly attenuated in ApoE^-/-^/Liver-DKO mice compared with ApoE^-/-^ controls, as well as in PCSK9-AAV8-induced Liver-DKO mice compared with PCSK9-AAV8-induced wild-type (WT) mice accompanied by reductions in blood cholesterol and triglyceride levels. Mechanistically, single-cell RNA sequencing of hepatocytes and aortas isolated from atherosclerotic ApoE^-/-^ and ApoE^-/-^/Liver-DKO mice revealed epsin-deficient Ldlr^hi^ hepatocytes with diminished lipogenic potential. Notably, pathway analysis of hepatocytes showed increased LDL particle clearance and enhanced LDLR-cholesterol interactions under WD treatment in ApoE^-/-^/Liver-DKO mice compared with ApoE^-/-^ controls, correlating with decreased plasma LDL-C levels. Furthermore, pathway analysis of the aortas showed attenuated inflammation and endothelial activation, coupled with reduced lipid uptake, and enhanced cholesterol efflux under WD treatment in ApoE^-/-^/Liver-DKO mice compared with ApoE^-/-^ controls. Moreover, the absence of liver epsins led to an upregulation of LDLR protein expression in hepatocytes. We further demonstrated that epsins bind LDLR via the ubiquitin-interacting motif (UIM), enabling PCSK9-mediated LDLR degradation. Depleting epsins abolished this degradation, thereby preventing atheroma progression. Lastly, targeting liver epsins with nanoparticle-encapsulated epsins siRNAs effectively ameliorates dyslipidemia and inhibits atherosclerosis progression. These results are consistent with findings showing an increased epsin1 and epsin2 expression in atherosclerotic cardiovascular disease patients.

**Conclusions:** Liver epsins drive atherogenesis by promoting PCSK9-mediated LDLR degradation, thereby elevating circulating LDL-C levels and heightening lesional inflammation. As such, targeting epsins in the liver represents a promising therapeutic strategy to mitigate atherosclerosis by preserving LDLR and enhancing LDL-C clearance in the liver.

## INTRODUCTION

Atherosclerosis is a primary driver of cardiovascular diseases (CVDs), including coronary artery disease, stroke, and peripheral artery disease^1^. CVDs remain the leading cause of mortality worldwide. In the United States alone, heart-related conditions account for approximately 610,000 deaths annually—nearly one-quarter of all deaths^2^. Atherosclerosis begins with low-density lipoprotein cholesterol (LDL-C) accumulation in the subendothelial space, triggering an inflammatory cascade that promotes plaque development and progression^3^. Elucidating the molecular mechanisms underlying plaque initiation, progression, and rupture is critical for identifying therapeutic targets and preventing ischemic events, disability, and death in patients with this chronic disease^4^.

Modern lifestyles have contributed to the growing global prevalence of hyperlipidemia, and elevated levels of LDL-C are an independent risk factor for atherosclerotic cardiovascular disease (ASCVD). Familial hypercholesterolemia (FH) represents the most common inherited metabolic disorder, characterized by markedly elevated plasma levels of LDL-C^5^ and increased risk of premature ASCVD. Mutations in the *Ldlr* gene, encoding the low-density lipoprotein receptor (LDLR), are the primary genetic drivers of LDL-C elevation in FH^6^. Accordingly, therapeutic strategies aimed at lowering LDL-C remain a cornerstone in the prevention and treatment of inflammation-driven atherogenesis.

The regulation of circulating LDL-C levels is in large part mediated through its clearance by low-density lipoprotein receptors (LDLRs) that are expressed on hepatocytes. LDLR is a transmembrane protein that facilitates the uptake of circulating LDL-C, serving as the primary mechanism for plasma LDL-C clearance^7^. Proprotein convertase subtilisin/kexin type 9 (PCSK9), also primarily produced by the liver, binds to LDLRs and promotes their lysosomal degradation, thereby reducing receptor recycling and increasing plasma LDL-C levels^8^. Consequently, PCSK9 has emerged as a key therapeutic target. PCSK9 inhibitors, which prevent the interaction between PCSK9 and LDLR, have demonstrated potent lipid-lowering effects in clinical trials and real-world use^9,10^.

Animal models deficient in LDLR, such as LDLR knockout (KO) mice and rabbits, have been instrumental in advancing the understanding of hyperlipidemia and atherosclerosis^11,12^. Furthermore, Keeter et al. demonstrated that liver-specific overexpression of a variant of PCSK9 via adeno-associated virus serotype 8 (AAV8) induces hyperlipidemia and accelerates atherosclerotic plaque formation^13^. More recently, a CRISPR-based gene-editing strategy—VERVE-101—enabled potent and durable inactivation of hepatic PCSK9 expression in nonhuman primates, resulting in substantial and sustained reductions in plasma LDL-C levels^14^. These findings underscore the therapeutic potential of enhancing LDL-C clearance by inhibiting PCSK9-mediated LDLR degradation as a highly effective strategy for treating hyperlipidemia.

Despite significant therapeutic advances, the molecular mechanisms underlying PCSK9-mediated LDLR degradation remain poorly understood. In particular, it is unclear whether specific endocytic adaptor proteins facilitate LDLR internalization and degradation in response to PCSK9. Elucidating these molecular processes may uncover novel therapeutic targets, especially for patients who develop resistance or exhibit limited responsiveness to current PCSK9 inhibitors.

Epsins, a family of endocytic adaptor proteins, have recently emerged as critical regulators and potential therapeutic targets of atherosclerosis^15^. We previously demonstrated that epsins 1 and 2 are upregulated in atherosclerotic plaques of apolipoprotein E-deficient (ApoE^-/-^) mice fed a Western diet (WD)^4,16,17^.

Endothelial cell (EC)-specific deletion of epsins markedly attenuated atherogenesis in this model, primarily by reducing arterial inflammation, downregulating adhesion molecule expression, limiting monocyte recruitment, and alleviating endoplasmic reticulum (ER) stress through stabilization of inositol 1,4,5-trisphosphate receptor type 1 (IP3R1) in ECs^4^.

Moreover, we demonstrated that myeloid-specific epsin deficiency in ApoE^-/-^ mice impeded lesion progression by inhibiting foam cell formation and enhancing efferocytosis within the plaque^16,17^. These mice also exhibited improved reverse cholesterol transport and promoted atheroma regression through the stabilization of ATP-binding cassette transporter G1 (ABCG1) in macrophages^16,17^. Despite the central role of epsins in the initiation and progression of atherosclerosis as well as in plaque regression, their potential involvement in hepatic regulation of circulating LDL-C levels via modulation of LDLR stability remains largely unexplored. Addressing this gap may provide mechanistic insights into the biological functions of epsin and unveil a novel therapeutic avenue for the treatment of hyperlipidemia and atherosclerosis.

To investigate the physiological role of hepatic epsins in regulating LDLR stability during atherosclerosis, we generated a novel ApoE^-/-^ mouse with liver-specific deletion of epsins 1 and 2 (ApoE^-/-^/Liver-DKO). Here, we demonstrated that these mice exhibit reduced plaque formation compared with ApoE^-/-^ mice on WD. We observed enhanced LDLR proteins in the liver of ApoE^-/-^/Liver-DKO compared with ApoE^-/-^ mice, suggesting that epsins are required for LDLR degradation, and that their loss enhances LDLR stability by preventing its lysosomal degradation. Mechanistically, epsins directly bind LDLR through their ubiquitin-interacting motif (UIM), and PCSK9-induced LDLR degradation is largely abolished in the absence of hepatic epsins.

Single-cell RNA sequencing analysis of aorta and liver revealed upregulation of pathways involved in LDL particle clearance in ApoE^-/-^/Liver-DKO mice, consistent with elevated hepatic LDLR expression. Consistently, in mice harboring the gain-of-function human *PCSK9* D374Y mutation, we observed elevated hepatic epsin expression and suppressed LDLR levels. When we introduced *PCSK9* D374Y in WT and Liver-DKO mice via AAV8, we observed striking inhibition of plaque development in Liver-DKO harboring *PCSK9* D374Y transgene—supporting a model in which liver epsins mediate PCSK9-driven LDLR degradation and promote atherogenesis. Furthermore, silencing hepatic epsins via delivery of nanoparticle-encapsulated epsins siRNAs effectively reduced dyslipidemia and atherosclerosis, establishing liver epsins as a promising therapeutic target for hyperlipidemia and cardiovascular disease.

## MATERIALS and METHODS

### Animal Models

In this study, all animal procedures were performed in compliance with institutional guidelines and mouse protocols were approved by the Institutional Animal Care and Use Committee (IACUC) of Boston Children’s Hospital, MA, USA. Both male and female mice were used. C57BL/6 mice (stock #00664), ApoE ^-/-^ mice (stock #002052), and Alb-Cre delete mice (stock #035593) were all purchased from Jackson Research Laboratory. As double knockout mice of Epsin 1 and 2 (Epsin 1^-/-^; Epsin2 ^-/-^) led to embryonic lethality, we generated conditional Epsin1 ^fl/fl^; Epsin2 ^-/-^ mice previously described^18^. ApoE ^-/-^ mice, Alb-Cre^+/-^ mice and Epsin1 ^fl/fl^; Epsin2 ^-/-^ mice were backcrossed to C57BL/6 background. The Alb-Cre^+/-^ mice were used for deletion of *loxP*-flanked Epsin1 in the liver. We bred Epsin1 ^fl/fl^; Epsin2 ^-/-^ mice with Alb-Cre^+/-^ mice to generate Epsin1 ^fl/fl^; Epsin2 ^-/-^; Alb-Cre^+/-^ hepatocyte-specific Epsins deficient (Liver-DKO) mice (Figure S1A). In addition, we bred Epsin1 ^fl/fl^; Epsin2 ^-/-^; Alb-Cre^+/-^ mice with ApoE^-/-^ (C57BL/6) background to generate Epsin1 ^fl/fl^; Epsin2 ^-/-^; Alb-Cre^+/-^; ApoE ^-/-^ mice (ApoE ^-/-^/Liver-DKO) (Figure S1B).

The control mice for Epsin1 ^fl/fl^; Epsin2 ^-/-^; Alb-Cre^+/-^ (Liver-DKO) mice were Epsin ^+/+^; Epsin2 ^+/+^; Alb-Cre ^+/-^ mice (WT) (Figure S1A). Western blot and immunofluorescence analyses of Epsins 1 and 2 in hepatocytes isolated from Liver-DKO versus WT mice confirmed that both Epsin1 and Epsin2 were markedly downregulated in livers of Liver-DKO mice (Figures S2A and S2B). The control mice for Epsin1 ^fl/fl^; Epsin2 ^-/-^; Alb-Cre^+/-^; ApoE ^-/-^ mice (ApoE^-/-^/Liver-DKO) were Epsin1 ^+/+^; Epsin2 ^+/+^; Alb-Cre ^+/-^; ApoE^-/-^ (ApoE^-/-^) (Figure S1B). For control mice, in addition to ApoE^-/-^; Epsin 1^+/+^; Epsin 2 ^+/+^ mice, we also used ApoE^-/-^; Epsin 1^+/+^; Epsin 2 ^+/+^ mice with a single copy of Alb-Cre, and ApoE ^-/-^ mice; Epsin1 ^fl/fl^; Epsin2 ^-/-^ littermates lacking the single copy of Alb-Cre. To simplify the terminology, we refer to these control mice as ApoE^-/-^, as results were not different in any of the analyses we performed.

To induce atherosclerosis, mice were fed a Western diet (WD, Protein 17% kcal, Fat 40% kcal, Carbohydrate 43% kcal; D12079B, Research Diets, New Brunswick, USA) starting at the age of 6–8 weeks for 8–20 weeks. Mice were sacrificed at different time points based on experimental design and the liver, blood, heart, and aorta were harvested. For each experimental model and time point, 6–10 mice were analyzed and both male and female mice were used in separate groups. In the current study, we did not exclude any mice when analyzing the data.

### Liver and Aorta Single-cell Preparation and Single-cell RNA Sequencing

For liver cell isolation, the ApoE^-/-^ and ApoE^-/-^/Liver-DKO mice (n=3) were anesthetized and restrained, and the skin sprayed with 70% ethanol. The liver and other inner organs were revealed by cutting through the skin and peritoneum. A 24G needle was carefully inserted into the inferior vena cava and secured with a clamp, and chelating solution (0.05M 4-(2-hydroxyethyl)-1-piperazineethanesulfonic acid (HEPES), pH 7.2, 10 mM ethylene glycol-bis(β-aminoethyl ether)-N, N, N’, N’-tetraacetic acid (EGTA) in Hank’s balanced salt solution (HBSS) without CaCl_2_ and MgCl_2_) was run at a low speed (1.5–2 mL/minute). The portal vein was then cut, and perfusion speed was increased to a flow rate of 7 mL/minute. After that, the diaphragm was cut and the anterior vena cava clamped. The chelating perfusion was run for 7 minutes and then switched to collagenase solution (0.05 M HEPES, pH 7.2, 4.7 mM CaCl_2_, 20 μg/mL Liberase, Sigma LIBTM-RO) at a flow rate of 7 mL/minute for 7 minutes. The liver was then removed and passed through a 70 μm cell strainer with 10 ml ice-cold HBSS without CaCl_2_ and MgCl_2_. The resulting single-cell suspension was centrifuged at 300 g for 5 minutes at 4 ℃ and washed ice-cold HBSS. The liver cell pellet was suspended in phosphate-buffered saline (PBS) buffer containing 0.04% bovine serum albumin (BSA).

For the aortic sample preparation, we used our previously reported protocol with minor modification^19^. Briefly, mouse aortas isolated from the ApoE^-/-^ and ApoE^-/-^/Liver-DKO mice (n=3) were cut into small pieces and placed in 6.5 mL of enzyme solution at 37 ℃ for 1.5 hours (5 mL of Dulbecco’s Modified Eagle Medium (DMEM), 1 mL of trypsin, 10 mg/mL collagenase type I, 10 mg/mL collagenase type IV, 50 μL elastin [stock:2.5 mg/mL], 200 μL of collagenase type II [stock: 1 mg/mL], 200 μL of collagenase XI [stock: 1 mg/mL], 200 μL of hydrolase [stock: 1 mg/mL], 100 μL of Liberase [stock: 3.85 mg/mL, Roche], 10 μL of DNase I [stock: 5 U/ μL, Sigma]). This solution was then filtered with a 40 μm cell strainer, centrifuged (400 g for 5 minutes), and the cell pellet was suspended in a PBS buffer containing 0.04% BSA. The isolated liver and aorta cells were counted and diluted into 1000 cells/ μL. The single-cell RNA-seq library construction was performed according to the 10x genomics protocol (GEMs, Single cell 3’ Reagent Kits v3.1, 10x Genomics). The prepared libraries were sequenced on the HiSeq 2500 Sequencing System (Illumina, San Diego, CA) with 100-bp paired end reads.

### Single-cell Transcriptome Data Analysis

First, we employed Cell Ranger (version 7.1.0) to map the raw reads of RNA sequencing data to the mouse genome (version mm10) and count the unique molecular identifier (UMI) for each gene. We then proceeded with the resulting UMI count matrix using the Seurat R package (version 4.3.0)^20^. We retained high-quality cells expressing between 200 and 2500 genes, excluding those with over 20% mitochondrial reads in the liver datasets and 5% mitochondrial reads in the aorta datasets. Additionally, we filtered out rarely expressed genes detected in fewer than 3 cells. After filtering, the high-quality data was normalized, cell-level scaled with 10k reads, and natural-log transformed by adding 1. The normalized data underwent further processing steps: scaling (ScaleData function), principal component analysis (PCA) (RunPCA function, npcs =30), Uniform Manifold Approximation and Projection (UMAP) (RunUMAP function, reduction = “pca”, dims = 1:30), shared nearest neighbor graph (SNNG) construction (FindNeighbors function, reduction = “pca”, dims = 1:30), and cell clustering (FindCluster function, resolution=0.1).

We conducted differentially expressed gene (DEG) analyses in one cluster versus other clusters using the FindAllMarkers function. The Wilcoxon test method was used by default, with a minimum percentage of expressed cells set to 25% and a minimum log2 fold change of 0.25. Cell types were annotated based on known marker genes from PanglaoDB^21^, cell-Taxonomy^22^, disco^23^ databases, and relevant literature. Marker gene expressions were visualized by DotPlot and VlnPlot functions. For cell rank for directed single-cell transition mapping, we utilized cellrank (version 2.0.4)^24^ with default parameters. The metabolite-mediated cell-cell communication was analyzed by MEBOCOST (version 1.0.0)^25^. The data were analyzed combined for both conditions following the tutorial on the MEBOCOST website (https://github.com/zhengrongbin/MEBOCOST). The prediction of sender-metabolite-sensor-receiver communication events was visualized by the bar and lollypop plots, using the ggplot2 library. Additionally, we performed cell-cell communication analysis using Cellchat (version 1.5.0)^26^. The communication probability of each condition was analyzed to highlight differences between conditions, and communication events were visualized using bar, flow, and circle plots. Furthermore, Gene Ontology (GO) functional enrichment analysis was performed using the clusterProfiler R package (version 3.18.1)^27^, and visualized by bar, lollipop, and cnet plots.

Finally, our research involved a comprehensive approach to quantifying the cardioprotective potential based on gene expression. We began by compiling a curated list of genes associated with the clearance of low-density lipoprotein cholesterol (LDL-C) in the liver that were mapped to the risk genes of coronary artery disease (CAD) identified by a genome-wide association study (GWAS), such as *Ldlr*, *Apoe*^28^. The risk genes for coronary artery disease are listed in Table S1. We then computed a CAD protective score for each sample. First, we extracted normalized expression values of the selected CAD protective genes. This process involved collecting data on the expression levels of these genes from each sample, which was then normalized to account for any variations in the data collection process. The CAD protective score was then calculated as the average expression of these genes chosen across each sample, providing a single composite metric representing overall protective gene activity. We employed violin plots to evaluate differences in CAD protective scores between experimental groups. These plots visualized the distribution of scores across conditions, allowing both the spread and central tendency to be assessed. The statistical significance of these differences was then rigorously evaluated using a t-test, a crucial step in our analysis. In addition to data analysis from ApoE^-/-^ and ApoE^-/-^/Liver-DKO mice, we reanalyzed the scRNA-seq data from PCSK9 D374Y mutant with C57BL/6 genetic background (GSE254971) and compared with our healthy control C57BL/6 mice using the methods described above.

### Human Samples

The healthy controls and diseased aortic arch samples from atherosclerosis patients were purchased from Maine Medical Center Biobank. The medical information details of the atherosclerotic patients and healthy people samples are listed in Table S1. The paraffin sections were deparaffinized, and we performed antigen retrieval to unmask the antigenic epitope with 10 mM Sodium Citrate, pH 6.0, with 0.5% Tween 20 at 95 °C for 30 minutes. Then the slides were allowed to cool for 30 minutes before staining.

### Oil Red O Staining

Whole aortas were dissected symmetrically, pinned to parafilm to allow the *en face* to be exposed and fixed in formalin for 12 hours. Aortas were washed in PBS 3 times and rinsed in 100% propylene glycol followed by staining with 0.5% Oil Red O solution for 20 minutes at 65 °C. Aortas were then put in 85% propylene glycol for 2 minutes, followed by three washes in DD Water. Imaging was performed using a Nikon SMZ1500 stereomicroscope, SPOT Insight 2Mp Firewire digital camera, and SPOT Software 5.1. The *en-face* Oil Red O-stained areas were manually traced and further quantified by NIH ImageJ software with n=3 or more. Hearts were embedded in O.C.T compound and sectioned at 8 microns. Cryostat sections of mouse aortic root were fixed in 4% paraformaldehyde for 15 minutes. Slices were washed in PBS followed by staining with freshly prepared 0.5% Oil Red O solution in isopropanol for 10 minutes at 37 °C. Slices were then put in 60% isopropanol for 30 seconds, followed by 3 washes in water. Slices were mounted with 90% Glycerin. The Oil Red O (ORO)-stained aortic root lesion areas were quantified by NIH ImageJ software with n=3 or more.

### Periodic Acid-Schiff’s Staining

Periodic acid-Schiff’s staining was performed according to manufacturer’s instructions. In brief, paraffin sections of mouse liver samples were deparaffinized by xylene (2 changes, each for 5 minutes) and followed by rinsing in absolute alcohol for 3 changes (each for 1 minute) and were rehydrated by running distilled water for 1 minute. The slides were immersed in 0.5% Periodic acid for 7 minutes and followed by rinsing in running distilled water for 10 seconds. The slides were further immersed into Optimized Schiff’s Water for 15 minutes and followed by rinsing in running tap water for 5 minutes. For nuclear staining, the slides were immersed in Modified Mayer’s Hematoxylin for 2 minutes and then rinsed in running tap water for 1 minute. The slides were further immersed into Light Green to counterstain until the desired background intensity is reached. The stained slides were dehydrated by absolute alcohol (3 changes, each for 1 minute) and cleared by xylene (3 changes, each for 1 minute) and mounted using Permount (ThermoFisher). The quantification of glycogen-stained areas in the liver samples was performed by NIH ImageJ software with n=3 or more.

### Immunofluorescence Staining

The liver samples and aortas from both ApoE^-/-^ and ApoE^-/-^/Liver-DKO mice were subjected for cryosections, and sections were further fixed in 4% paraformaldehyde for 15 minutes. The slides were blocked by blocking buffer (PBS/3% BSA/3% donkey serum/0.3% triton) for 30 minutes, and were further incubated by primary antibodies (antibody list is shown in Major Resource Table) for overnight at 4°C. The slides were washed 3 times for 10 minutes each wash in PBS/0.3% triton buffer and were then incubated with secondary antibodies at room temperature for 1 hour. Next, the slides were washed 3 times for 10 minutes, each washed in PBS/0.3% triton buffer. After the second wash, DAPI was used for nuclei staining. The slides were mounted using a Fluoroshield mounting medium. Imaging was performed using Zeiss LSM 880 Confocal Acquisition & Analysis. Samples stained without the primary antibody were obtained using the same settings as negative controls (Figure S14). Mean fluorescence intensity (MFI) of antibody staining was determined using NIH ImageJ software with n=3 or more.

### Synthesis of siRNA Nanoparticles (NPs)

In this study, a robust self-assembly method was used to prepare the polymer-lipid hybrid NPs for siRNA delivery^29^. In brief, G0-C14 and PLGA were dissolved separately in anhydrous dimethylformamide (DMF) to form a homogeneous solution at concentrations of 2.5 mg/mL and 5 mg/mL, respectively. DSPE-PEG-OCH_3_ (DSPE-mPEG) was dissolved in HyPure water (GE Healthcare Life Sciences, catalog no. SH30538) at a concentration of 0.1 mg/mL. 1 nmole Epsin1 siRNA and 1 nmole Epsin2 siRNA were gently mixed with 100 μL of the G0-C14 solution. The mixture of siRNA and G0-C14 was incubated at room temperature for 15 minutes to ensure the full electrostatic complexation. Next, 500 μL of PLGA polymers were added and mixed gently. The resultant solution was subsequently added dropwise into 10 mL of HyPure water containing 1 mg DSPE-mPEG under magnetic stirring (1,000 rpm) for 30 minutes. The siRNA NPs were purified by an ultrafiltration device (EMD Millipore, MWCO 100 kDa) to remove the organic solvent and free excess compounds via centrifugation at 4 °C. After washing 3 times with HyPure water, the siRNA NPs were collected and finally resuspended in pH 7.4 PBS buffer. The NPs were used freshly or stored at -80 °C for further use.

### RNA Isolation and Real-time Quantitative PCR

Total RNA was extracted from the liver tissue with the RNeasy Mini Kit (Qiagen) according to the manufacturer’s instructions, including the optional step to eliminate genomic DNA. The extracted RNA was used for RT-qPCR according to the experimental designs. For the RT-qPCR, mRNA was reverse-transcribed to cDNA with the iScript cDNA Synthesis Kit (Bio-Rad Laboratories). 2 μL of 5-fold diluted cDNA product was subjected to the RT-qPCR in StepOnePlus Real-Time PCR System (Applied Biosystems) using SYBR Green PCR Master Mix reagent as the detector. PCR amplification was performed in triplicate on 96-well optical reaction plates and replicated in at least three independent experiments. The Delta-Delta Ct (ΔΔCt) method was used to analyze qPCR data. The Ct of β-actin cDNA was used to normalize all samples. Primers are listed in the Major Resource Table.

### Analysis of Plasma Cholesterol and Triglyceride Levels

Blood was removed from the right atrium of the mouse heart after sacrifice with isoflurane. Blood was allowed to clot for 30 minutes at room temperature followed by centrifugation at 3000 g at 4 °C for 20 minutes. Serum was transferred to a new tube and stored at -20 °C. Serum cholesterol and lipid levels were determined using the Cholesterol Assay Kit and Triglyceride Assay Kit (Abcam).

### Immunoprecipitation and Western Blotting

For immunoprecipitation, HepG2 cells were lysed with radioimmunoprecipitation assay (RIPA) buffer (50 mM Tris, pH 7.4, with 150 mM NaCl, 1% Nonidet P-40, 0.1% sodium dodecyl sulfate (SDS), 0.5% sodium deoxycholic acid, 0.1% sodium dodecyl sulfate, 5 mM N-ethylmaleimide and protease inhibitor cocktail). For LDLR ubiquitination experiments, HepG2 cells were lysed using a denaturing buffer (1% SDS in 50 mM Tris, pH 7.4) and boiled at 95 °C for 10 minutes to denature protein complexes. Lysates were re-natured using nine volumes of ice-cold RIPA buffer, then prepared for immunoprecipitation as follows. Cell lysates were pre-treated with Protein A/G Sepharose beads at 4 °C for 2 hours to remove nonspecific protein followed by centrifugation at 12000 rpm for 5 minutes at 4 °C. Supernatant was transferred to a new tube, incubated with Protein A/G Sepharose beads and antibodies against Epsin1 or LDLR or ubiquitin at 4 °C overnight. Mouse IgG was used as a negative control. Protein A/G beads were washed with RIPA buffer 2 times, followed by PBS once. Then, beads were suspended with an 80 μL 2x loading buffer and heated at 95 °C for 10 minutes. After centrifugation, precipitated proteins were visualized by Western blot. Proteins were resolved by SDS-PAGE gel and electroblotted to nitrocellulose membranes. NC membranes were blocked with 5% milk (w/v) and blotted with antibodies. Western blots were quantified using NIH Image J software with n=3 or more. The raw films for western blots exposure were submitted in Supplemental Material.

### PCSK9 Adeno-associated Virus 8 (AAV8) Tail Vein Injection

Eight-week-old male C57BL/6 mice, both WT and Liver-DKO mice (n=6), were intravenously injected via tail vein with a single dose of 2 x 10^11^ viral PCSK9-AAV8 and fed a WD for 20 weeks. The serum samples collected from both WT and Liver-DKO mice were subjected for triglyceride and cholesterol measurements. The aortas were isolated for *en face* ORO staining and histology analysis, and the liver tissue from both WT and Liver-DKO mice was collected for further histology analysis and protein lysate preparation.

### Nanoparticle-encapsulated Epsin1 and Epsin2 siRNAs Tail Vein Injection

Eight-week-old male C57BL/6 mice, ApoE ^-/-^ mice were fed a WD for 8 weeks and further divided into two groups (n=6). The control group mice were intravenously injected via tail vein with control siRNA NPs, and the experimental group mice were injected with 1 nmoles^29^ epsin1 and epsin2 siRNA NPs for continuous three weeks. Two injections per week. After injection, the serum samples collected from both control siRNA and epsin 1 and 2 siRNA NPs injected groups were subjected to triglyceride and cholesterol measurement. The aortas were isolated *en face* ORO staining and histology analysis, and the liver tissue from both control siRNA and epsin 1 and 2 siRNA NPs were collected for protein lysate preparation.

### Cell Culture and Plasmids Transfection

The HepG2 cell line (ATCC no. HB-8065) was used for plasmid transfection to map the binding sites of Epsin1 to LDLR. Flag-tagged Epsin1WT, Epsin1ΔUIM, Epsin1ΔENTH, Epsin1-DPW/NPF truncation constructs, and pcDNA vector were prepared previously in our laboratory. HepG2 cells were cultured in DMEM (10% fetal bovine serum (FBS) and 1% Pen-Strep) at 37°C in humidified air containing 5% CO_2_ atmosphere and transfected using Lipofectamine 2000 as instructed by the manufacturer.

### Epsin1 and Epsin2 siRNAs Transfection, PCSK9-AAV8 Infection

HepG2 cells were cultured in DMEM (10% FBS and 1% Pen-Strep) at 37°C in humidified air containing 5% CO_2_ atmosphere. One day before transfection, plate cells in 1 mL of growth medium without antibiotics such that they will be 30-50% confluent at the time of transfection. The transfection of epsin1 and epsin2 siRNAs was performed according to the manufacturer’s instructions. The preparation of RNAi duplex-Lipofectamine RNAiMAX complexes by mixture of either control siRNA or epsin1 and epsin2 siRNAs (Dharmacon, Horizon Discovery) and Lipofectamine RNAiMAX, maintaining the mixture at room temperature for 20 minutes, and then adding the complexes to each well containing cells. The cells were incubated for 48 hours at 37°C in a CO_2_ incubator. The PCSK9-AAV8 stock (10^13^ GC/ml) was diluted by culture media into 10^10^ GC for the infection. The original cell culture media were removed and added to the PCSK9-AAV8 containing media for cell culture. The cells were collected 3 days after the PCSK9-AAV8 infection.

### Hepatocyte Primary Culture, MG132 Treatment, PCSK9-AAV8 Infection

The methods of perfusion and digestion of liver were reported in liver single-cell preparation part above. The digested liver was transferred to a 10 cm plate with plating media (DMEM low glucose, 5% FBS, 1% Penicillin-Streptomycin Solution), and the liver cells were gently released with fine tip forceps. The liver cell suspension was filtered through a 70 μm cell strainer into a 50 mL tube and then spun at 50 g for 2 minutes at 4 °C. While the samples were spinning, a fresh Percoll solution (90% Percoll in 1xHBSS) was prepared. The supernatant was aspirated, and 10 mL of plating media was added and resuspend by gentle swirling. Next, 10 mL Percoll solution was added and mixed thoroughly by inverting the tube several times. After spinning at 200 g for 10 minutes at 4 °C, the supernatant was aspirated, 20 mL of plating media was added, and then spun at 50 g for 2 minutes at 4 °C. The supernatant was aspirated, and 20 mL of plating media was added. Hepatocytes were counted and plated on collagen-coated cell culture 6-well plates. After 3 hours, the medium was changed to warm maintenance media (Williams E media, 1% Glutamine, 1% Penicillin-Streptomycin Solution). Three hours after plating, the proteasome inhibitor MG132 solution was added to the cultured primary hepatocytes. After 6 hours, the primary hepatocytes were subjected to PCSK9-AAV8 infection as mentioned above. The primary hepatocytes were collected and lysed for protein preparation.

### DiI-LDL Uptake Assay

Hepatocytes isolated from WT and Liver-DKO mice were seeded plated in black clear bottom 96-well plates and further subjected to PCSK9-AAV8 infection. During the last 5 hours of PCSK9-AAV8 infection, hepatocytes were exposed to DiI-LDL (10 μg/ml) and then washed with two changes of PBS. The intracellular fluorescence intensity of DiI-LDL was then quantified using the SpectraMax Gemini XPS/EM Microplate Readers (Molecular Devices; ex 554/em 571). In addition, isolated hepatocytes were seeded in plated 12-well plates with sterilized coverslips, and hepatocytes were subjected to PCSK9-AAV8 infection and were further exposed to DiI-LDL (10 μg/ml). The hepatocytes were washed with two changes of PBS and were further fixed by 4% PFA for 10 minutes. DAPI was used for nuclei staining after PBS washing. The coverslips were mounted using a Fluoroshield histology mounting medium (Sigma-Aldrich). Imaging was performed using Zeiss LSM 880 Confocal Acquisition & Analysis. Mean fluorescence intensity (MFI) of DiI-LDL fluorescence was determined using NIH ImageJ software with n=3 or more.

### Avoiding Bias in Quantitative Analysis

We had two blinded observers select representative images from a panel of images collated from all experiments performed on any given sample. Representative images of immunofluorescence, Oil Red O and Periodic acid-Schiff’s staining were selected based on the most accurate representation of similarity with the mean value for each experimental group. The way to select a representative image would be an image that is most similar to all the other images in the set. Representative images from immunofluorescence, Oil Red O and Periodic acid-Schiff’s stains were selected based on high quality, resolution, and accurate representation of similarity with the mean value of each experimental group.

### Statistical Analysis

Gene expression was assessed by quantifying mRNA levels of target genes via qPCR, with normalization to the internal control, β-actin. The images of western blotting, immunofluorescence, aortic lesion sizes, Oil Red O, and Periodic acid-Schiff’s-stained images were quantified by ImageJ software (NIH, Bethesda, MD) running on Apple OS X. Statistical analysis of samples including Oil Red O, IF, and Periodic acid-Schiff’s stains were performed by blinding in which each animal was assigned a number and data was collected based on the assigned number with genotype and experimental condition unknown to the data collector. Quantitative data were analyzed by 2-tailed, unpaired, or paired Student’s t-test or two-way ANOVA with post hoc test, using Prism 8.0 (GraphPad Software) running on Apple OS X. All data are presented as are the mean ± SEM.

### Data Availability

The liver and aorta scRNA-seq data (GSE273386) in ApoE^-/-^ and ApoE^-/-^ /Liver-DKO mice are available in the Gene Expression Omnibus. The scRNA-seq data (GSE254971) for D374Y mCherry-APOB mice are available in the published paper entitled “Kupffer cells dictate hepatic responses to the atherogenic dyslipidemic insult” (https://doi.org/10.1038/s44161-024-00448-6). Source data is provided with this paper.

## RESULTS

### Elevated Epsin1 and Epsin2 Expression in the Aortic Biopsies of Atherosclerotic Cardiovascular Disease Patients

We measured the protein expression of epsin1 and epsin2 between healthy controls and atherosclerotic cardiovascular disease (ASCVD) patients by immunofluorescence staining. Intriguingly, we found 98% and 68% increases of epsin1 and epsin2 protein levels, respectively, in diseased aortic samples (Figure S3A-S3C). As expected, we found remarkably elevated macrophage marker CD68 protein expression in the aortic biopsies of ASCVD patients (Figure S3A-S3C), indicating more macrophage accumulation and increased atherosclerotic lesions in ASCVD patients. Importantly, we discovered a substantially higher percentage of colocalization between epsin1 and CD68, and a higher overlay percentage between epsin2 and CD68 in the macrophages in ASCVD patients than in healthy controls (Figure S3A-S3C).

### Depletion of Epsins in the Liver Decreases Atherosclerotic Lesion, Macrophage Accumulation, and Plasma Lipid Levels

We explored the physiological role of hepatic epsins in lipoprotein metabolism and atherosclerosis by employing liver-specific epsins-deficient mice (ApoE^-/-^/Liver-DKO) and compared their phenotypic outcomes with ApoE^-/-^ controls. Notably, in ApoE ^-/-^/Liver-DKO mice fed a WD, plasma cholesterol and triglycerides were dramatically reduced by 55% and 34%, respectively vs ApoE ^-/-^ controls (Figure 1A). *En face* Oil Red O staining of aortas from ApoE ^-/-^ and ApoE ^-/-^/Liver-DKO mice fed a WD revealed a 69% reduction in atherosclerotic lesion area in ApoE^-/-^/Liver-DKO mice compared with ApoE ^-/-^ controls (Figure 1B). The lesion areas of the aortic roots in ApoE ^-/-^/Liver-DKO mice revealed a 67% reduction in mean atherosclerotic lesion area compared with ApoE ^-/-^ controls (Figure 1C). These findings indicate that liver epsins likely play crucial roles in lipoprotein metabolism with a major impact on the pathogenesis of atherosclerosis.

**Figure 1:**
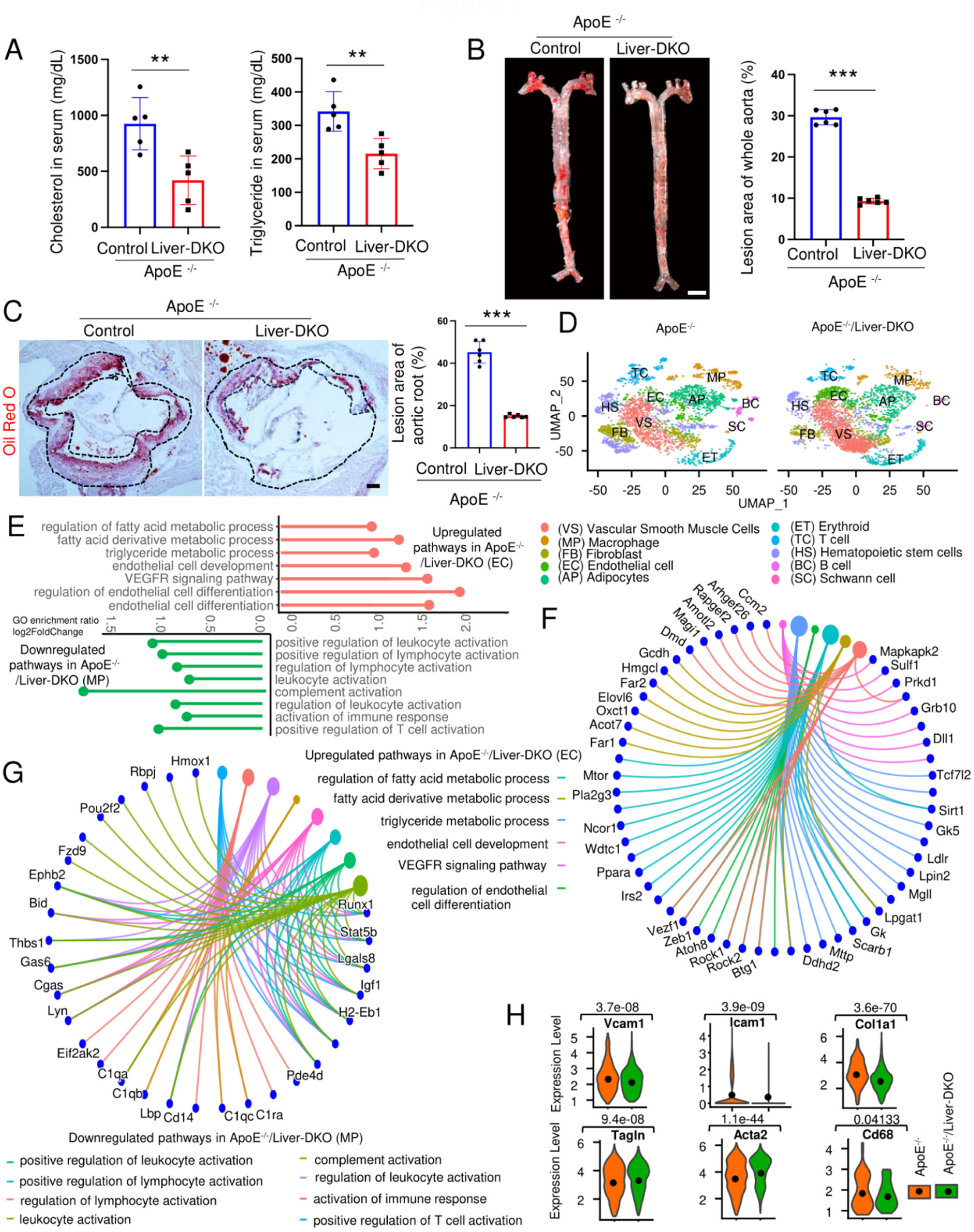
Liver-deficiency of epsins reduces plasma lipids and inhibits atherosclerotic lesion formation by altering the inflammatory cellular landscape of plaques in ApoE^-/-^ mice. A: Serum cholesterol and triglyceride (TG) levels in ApoE^-/-^ and ApoE^-/-^ /Liver-DKO mice after 8 weeks on a WD (n=5, p**<0.01). B: *En face* ORO staining of aortas (left) from ApoE^-/-^ or ApoE^-/-^/Liver-DKO mice fed a WD for 8 weeks, and unpaired t-test (right) for the lesion areas (n=6, ***p< 0.001). Scale bar, 5 mm. C: Aortic roots from WD-fed ApoE^-/-^ or ApoE^-/-^/Liver-DKO mice stained with ORO (left), and unpaired t-test (right) for the lesion areas (right) (n=6, ***p< 0.001). Scale bar, 200 μm. D: UMAP visualization illustrating the heterogeneity of cell populations indicating distinct clustering patterns of Macrophages (MP), Vascular Smooth Muscle Cells (VS), Endothelial Cells (EC) and Fibroblasts (FB) and other cells in ApoE^-/-^/Liver-DKO mice compared to ApoE^-/-^ controls. E: Gene Ontology (GO) analysis showing significantly enriched pathways for the regulation of triglyceride and fatty acid metabolic process, endothelial cell differentiation, vascular endothelial growth factor receptor (VEGFR) signaling pathways are upregulated in endothelial cells (EC) (top) and downregulated pathways of activation of inflammation in the macrophages (MP) (bottom) in ApoE^-/-^/Liver-DKO mice relative to ApoE^-/-^ controls. F: Circular cnetplot representing downregulated pathways and corresponding genes in macrophages (MP). G: Circular cnetplot illustrating significantly upregulated pathways and corresponding genes in endothelial cells (EC). H: Violin plots showing diminished expression of inflammation genes, such as CD68, Vcam1, Icam1, and downregulated expression of FB markers, such as Col1a1, and elevated expression of VS markers, such as Acta2 and Tagln. Statistical analysis (A, B, C, H) in ApoE^-/-^ and ApoE^-/-^/Liver-DKO comparison is conducted by the Student’s t-test.

### Reduced Inflammatory Macrophages, Attenuated Endothelial Inflammation, Enriched Contractile Vascular Smooth Muscle Cells, and Reduced Proportion of Fibroblasts in the Aortas in ApoE^-/-^/Liver-DKO Mice

We investigated cellular alterations in the aortas of ApoE^-/-^ and ApoE^-/-^/Liver-DKO mice by single-cell RNA sequencing (Figure S4A). The UMAP visualization uncovered 10 major cell types annotated according to the expression patterns of cell type-specific marker genes (Figure 1D, Figure S4B). Additional analyses revealed a reduced proportion of macrophages expressing CD68 in the aortas of ApoE^-/-^/Liver-DKO mice that had diminished expression of inflammatory pathway genes involving complement, leukocyte, and lymphocyte activation (Figure 1D, 1F, 1H, Figure S5D). In addition, the proportion of endothelial cells with no significant difference in expression of endothelial cell marker genes in the aortas in ApoE^-/-^/Liver-DKO mice, such as Pecam1/CD31, was increased (Figure 1D, Figure S4C, S4D). Importantly, the mRNA levels of the inflammatory monocytic adhesion molecules, including VCAM-1 and ICAM-1, were decreased in the endothelial layer of aortas in ApoE^-/-^/Liver-DKO versus ApoE^-/-^ mice (Figure 1H). Further analyses identified upregulated pathways involved in regulation of endothelial cell differentiation, endothelial cell development, vascular endothelial growth factor receptor signaling pathway, regulation of fatty acid metabolic process, and triglyceride metabolic process and downregulated pathways of regulation of endothelial cell apoptotic process, regulation of endothelial cell migration, and endothelial cell activation in endothelial cells in the aortas of ApoE^-/-^/Liver-DKO mice (Figure 1E, 1G, Figure S5A, S5C, Figure S6C, S6E). Intriguingly, the percentage of vascular smooth muscle cells (VSMCs) with elevated expression of contractile VSMC marker genes in the aortas in ApoE^-/-^/Liver-DKO mice, such as Acta2/α-SMA and Tagln/SM22, was increased (Figure 1D, 1H, Figure S4C, S4D). In contrast, the proportion of fibroblasts (FB) with diminished expression of fibroblasts marker genes in the aortas in ApoE^-/-^/Liver-DKO mice, such as Col1a1/Collagen I, was reduced (Figure 1D, 1H, Figure 2D, Figure S4C, S4D). In addition, pathways of cell-matrix adhesion, extracellular matrix organization, and inflammatory response in both fibroblasts and VSMCs in the aortas of ApoE^-/-^/Liver-DKO mice are downregulated (Figure S5B, S5D, S5E, Figure S6F, S6G). Correspondingly, the *Pecam1-*associated pathway was upregulated in endothelial cells, and collagen-associated pathways were downregulated in the aortas of ApoE^-/-^/Liver-DKO mice compared with ApoE^-/-^ controls (Figure S6A, S6B).

**Figure 2:**
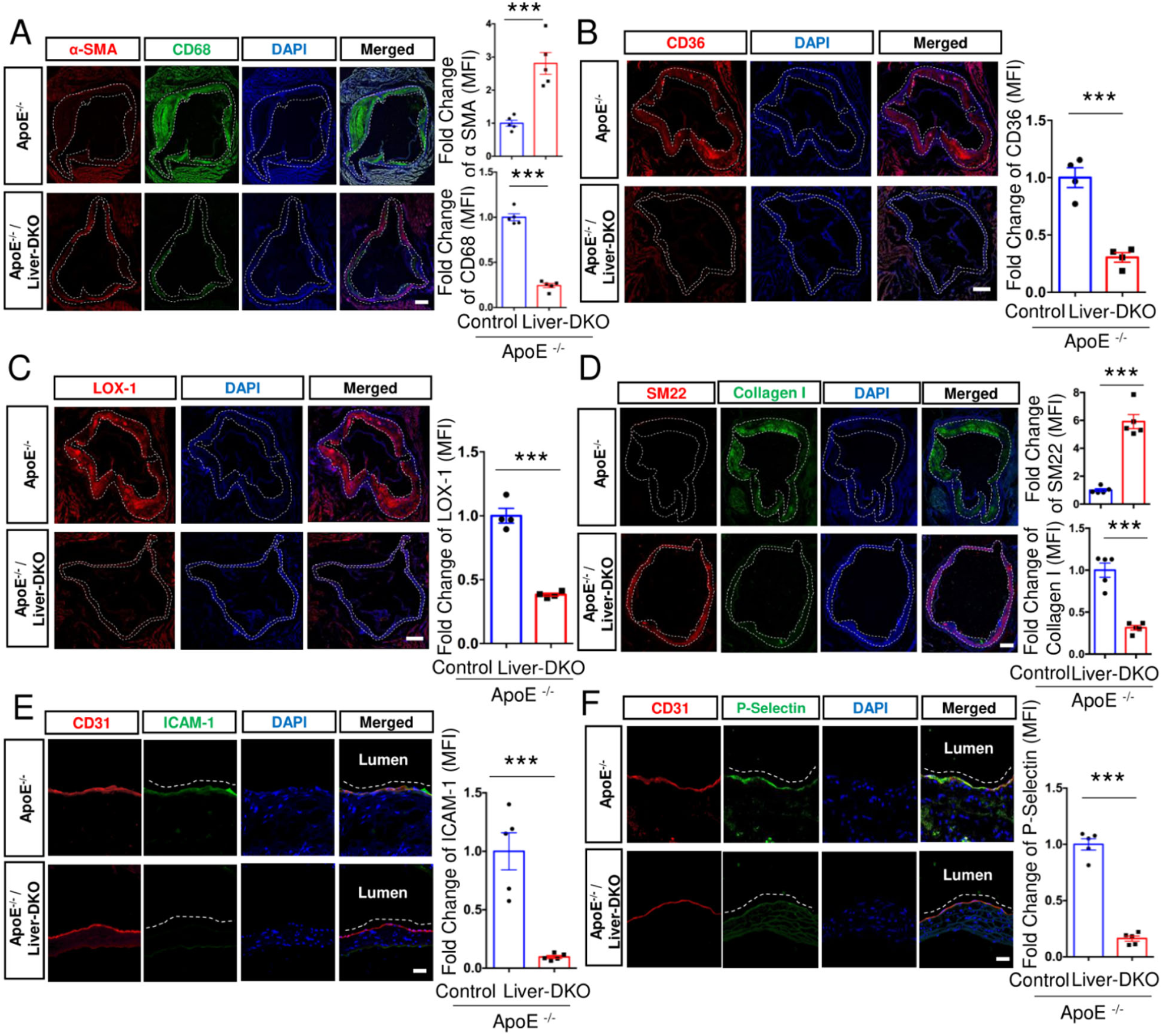
Liver-deficiency of epsins increases smooth muscle contractility, and inhibits inflammatory macrophage accumulation and endothelial cell activation in atherosclerotic plaques of ApoE^-/-^ mice. A: Immunofluorescence (IF) staining of vascular smooth muscle cell marker α-SMA and macrophage marker CD68 in the aortas of ApoE^-/-^ and ApoE^-/-^/Liver-DKO mice (left). The fold change of mean fluorescence intensity (MFI) for α-SMA and CD68 proteins were quantified (right) (n=5, ***p<0.001). Scale bar, 200 μm. Elevated expression of α-SMA protein and diminished expression of CD68 protein in the aortas in ApoE^-/-^/Liver-DKO. B-C: IF staining of scavenger receptors, such as CD36 (B) and LOX-1 (C), in the aortas of ApoE^-/-^ and ApoE^-/-^/Liver-DKO mice (left). The fold changes of CD36 (MFI) (B) and LOX-1 (MFI) (C) were quantified (right) (n=4, ***p<0.001). Scale bar, 200 μm. Diminished expression of CD36 and LOX-1 proteins in the aortas in ApoE^-/-^/Liver-DKO. D: IF staining of vascular smooth muscle cell marker SM22 and collagen marker Collagen I in the aortas of ApoE^-/-^ and ApoE^-/-^/Liver-DKO mice (left). The fold changes of SM22 (MFI) and Collagen I (MFI) were quantified (right) (n=5, ***p<0.001). Scale bar, 200 μm. Elevated expression of SM22 protein and diminished expression of Collagen I protein in the aortas in ApoE^-/-^/Liver-DKO. E-F: IF staining of endothelial cell marker CD31 and endothelial cell adhesion molecules, such as ICAM-1 (E) (left) and P-Selectin (F) (left), in the aortas between ApoE^-/-^ and ApoE^-/-^/Liver-DKO mice. The fold changes of ICAM-1 (MFI) (E) and P-Selectin (MFI) (F) were quantified (right) (n=5, ***p<0.001). Scale bar, 20 μm. Diminished expression of ICAM-1 and P-Selectin proteins in the aortas in ApoE^-/-^/Liver-DKO. All statistical analysis (A-F) in ApoE^-/-^ and ApoE^-/-^/Liver-DKO comparison is conducted by the Student’s t-test.

Consistent with the single-cell RNA sequencing analyses, we discovered remarkably diminished CD68 protein expression in the aortas in ApoE^-/-^/Liver-DKO mice compared with ApoE^-/-^ controls, demonstrating impeded macrophage accumulation in the atherosclerotic lesions (Figure 2A). Importantly, we detected attenuated inflammatory scavenger receptors including CD36 and LOX-1 in the aortas of ApoE^-/-^/Liver-DKO versus ApoE^-/-^ control mice (Figure 2B, 2C). In addition, the protein levels of α-SMA and SM22 were markedly increased in the aortas of ApoE^-/-^/Liver-DKO mice compared with ApoE^-/-^ controls which is consistent with increased contractile SMCs (Figure 2A, 2D). Furthermore, in keeping with the decreased number of fibroblasts in the aortas of ApoE^-/-^/Liver-DKO mice versus ApoE^-/-^ controls (Figure 1D), the collagen-1 protein levels were reduced (Figure 2D). Furthermore, the aortic endothelial layer in ApoE^-/-^/Liver-DKO versus ApoE^-/-^ mice had markedly decreased ICAM-1 and P-Selectin protein levels which is consistent with reduced endothelial cell activation and macrophage accumulation (Figures 2E, 2F).

### Depletion of Epsins in the Liver Negates the Effects of PCSK9 Overexpression on Atherosclerotic Lesion Formation, Macrophage Accumulation, and Plasma Lipid Levels

As overexpression of PCSK9 promotes atherosclerosis and elevates plasma lipids^13^, we further explored the role of hepatic epsins in lipoprotein metabolism and atherosclerosis by employing PCSK9-AAV8-induced Liver-DKO mice and compared their phenotypic outcomes with PCSK9-AAV8-injected WT controls. Mean plasma cholesterol and triglyceride (TG) levels showed striking 66% and 63% decreases resulting from loss of hepatic epsins in mice injected with PCSK9-AAV8 (2x10^11^ genomes) and fed a WD for 20 weeks (Figure S7A). In addition, low-density lipoprotein cholesterol (LDL-C) levels were reduced by 75% in 20 weeks WD-fed Liver-DKO mice after PCSK9-AAV8 injection (Figure S7A). Importantly, PCSK9-AAV8-induced Liver-DKO mice exhibited a 91% reduction in *en face* aortic lesion areas and a 92% reduction in lesions of the aortic roots compared with WT controls injected with PCSK9-AAV8 after 20 weeks WD treatment (Figure S7B, S7C). In addition, we found markedly attenuated CD68 expression in the aortas in PCSK9-AAV8-injected Liver-DKO mice compared with PCSK9-AAV8-injected WT controls (Figure S7D). Furthermore, the protein levels of α-SMA were markedly increased in the aortas of in PCSK9-AAV8-injected Liver-DKO mice versus PCSK9-AAV8-injected WT controls (Figure S7D). Importantly, the protein levels of the inflammatory endothelial cell adhesion molecules, such as ICAM-1 and P-Selectin, in the endothelial layer of aortas in PCSK9-AAV8-induced Liver-DKO mice (Figure S7E, S7F).

### Effects of Hepatic Deletion of Epsins 1 and 2 on Alb^hi^ and Ldlr^hi^ Hepatocyte Subpopulations and Lipoprotein Clearance Gene Expression

We show that deletion of Epsins1/2 markedly decreased plasma cholesterol and triglyceride. Therefore, we performed single-cell RNA sequencing on liver tissue derived from WD-fed ApoE^-/-^ and ApoE^-/-^/Liver-DKO mice to investigate the underlying mechanisms of the observed phenotypic changes (Figure S8A). The UMAP visualization revealed six different cell types based on the expression patterns of cell type-specific marker genes in the livers (Figure S8B, S8C). Analysis of the hepatocyte data revealed four distinct hepatocyte populations (Figure 3A, 3B, Figures S8B-8D), including HC1 Alb^hi^, HC2 Alb^hi^, HC3 Alb^hi^ and HC Ldlr^hi^, which are characterized by the expression levels of both *Alb* and *Ldlr* genes in hepatocytes. The expression of *Ldlr* gene is elevated from HC1 Alb^hi^ towards HC3 Alb^hi^, while with their lower *Ldlr* expression level compared with HC Ldlr^hi^. Notably, the proportion of Ldlr^hi^ hepatocytes increased by 59%, and the total proportions of HC2 Alb^hi^ and HC3 Alb^hi^ hepatocytes decreased by 25% and 20%, respectively, in ApoE^-/-^/Liver-DKO mice compared with the ApoE^-/-^ controls (Figure 3B).

**Figure 3:**
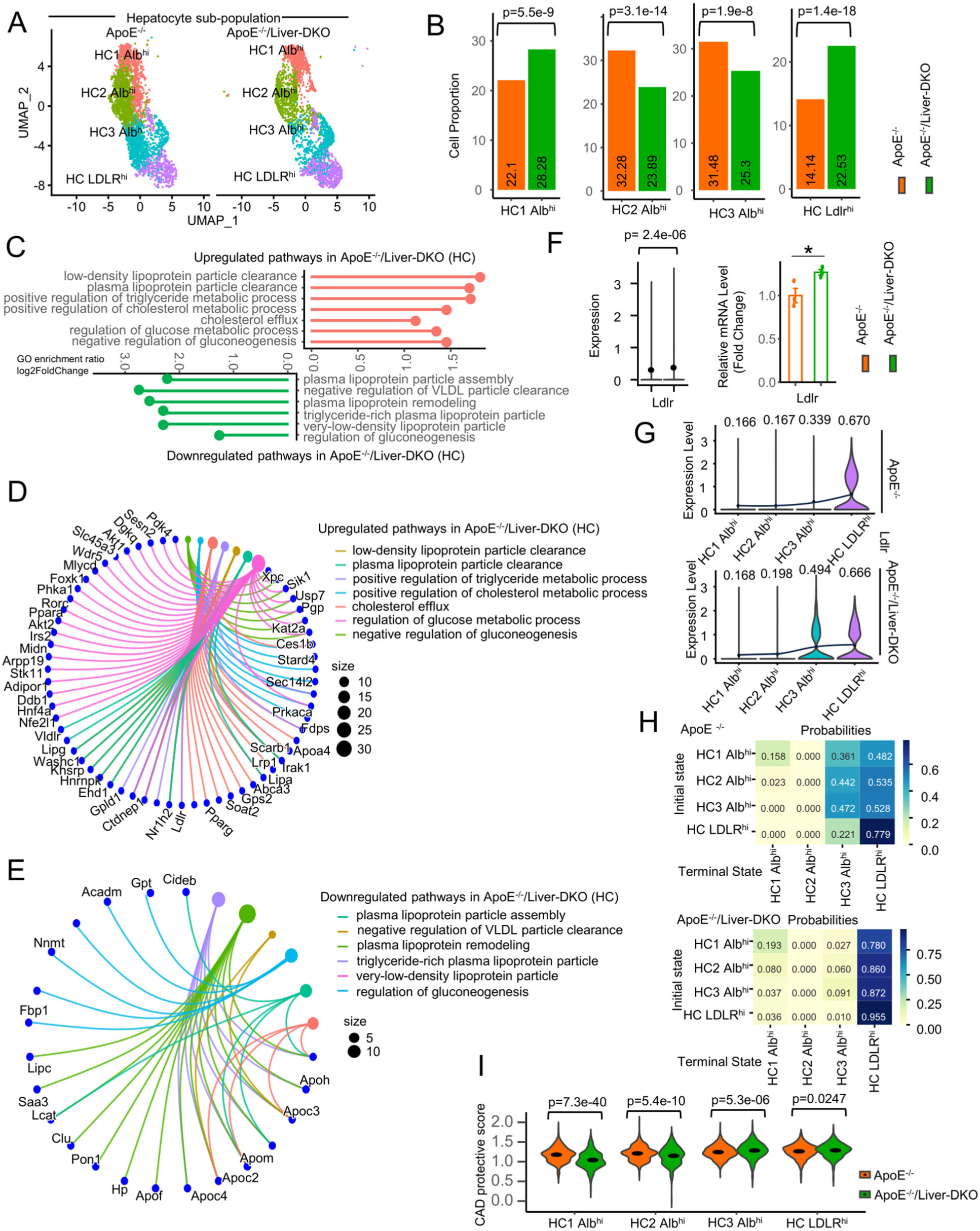
Single-cell RNA sequencing reveals increased Ldlr^hi^ hepatocytes proportion and elevated LDLR expression in the livers in ApoE^-/-^ /Liver-DKO mice compared to ApoE^-/-^ controls. A: UMAP visualization illustrating the heterogeneity of hepatocyte cell populations, indicating distinct clustering patterns of hepatocytes in ApoE^-/-^/Liver-DKO mice compared to ApoE^-/-^ controls. B: Bar plots of cell proportion of hepatocyte subclusters. There was a relatively high proportion of HC Ldlr^hi^ hepatocytes in the livers of ApoE^-/-^/Liver-DKO mice compared with ApoE^-/-^ controls. C: Gene Ontology (GO) analysis showing significantly enriched pathways of regulation of low-density lipoprotein particle clearance signaling pathways are upregulated in hepatocytes (HC) (top) and downregulated pathways of lipoprotein assembly and lipogenesis in hepatocytes (HC) (bottom) in ApoE^-/-^/Liver-DKO mice relative to ApoE^-/-^ controls. D: Circular cnetplot illustrating significantly upregulated pathways and corresponding genes in hepatocytes (HC). E: Circular cnetplot representing downregulated pathways and corresponding genes in hepatocytes (HC). F: Elevated expression of Ldlr in ApoE^-/-^/Liver-DKO mice compared with ApoE^-/-^ controls, shown through single-cell analysis (left) and real-time quantitative PCR (qPCR) (right). G: Violin plots of Ldlr expression in different hepatocyte subtypes, showing its upregulated expression in hepatocytes in ApoE^-/-^/Liver-DKO mice compared to ApoE^-/-^ controls. The mean expression levels of Ldlr are highlighted in the violin plots (p<0.05). H: CellRank analysis illustrating an increased propensity of differentiation from HC Alb ^hi^ towards HC Ldlr ^hi^ in ApoE^-/-^/Liver-DKO mice relative to ApoE^-/-^ controls. I: Coronary artery disease (CAD) protective score comparing ApoE^-/-^/Liver-DKO with ApoE^-/-^, showing it positively correlated to the expression of Ldlr (n=3, *p<0.05). Statistical analysis (B, F, G, I) in ApoE^-/-^ and ApoE^-/-^/Liver-DKO comparison is conducted by the Student’s t-test.

Hepatocyte-derived data analysis revealed upregulated pathways or genes of LDL particle clearance and downregulated pathways genes of hepatic lipogenesis and plasma lipoprotein particle assembly in different types of hepatocytes in ApoE^-/-^/Liver-DKO mice compared with the ApoE^-/-^ controls (Figure 3C-3E), e.g., *Ldlr*, suggesting improved LDL-C clearance in the absence of hepatic epsins. Consistent with the increased number of Ldlr^hi^ hepatocytes, there was overall elevated Ldlr expression in the livers from ApoE^-/-^/Liver-DKO versus ApoE^-/-^ mice as measured by both single-cell analysis and real-time quantitative PCR (Figure 3F). The hepatocyte subpopulations, particularly the Ldlr^hi^ and HC3 Alb^hi^ hepatocytes exhibited increased Ldlr expression (Figure 3G). Intriguingly, RNA velocity analysis supported an increased propensity for Alb^hi^ hepatocyte differentiation towards Ldlr^hi^ hepatocytes in the absence of hepatic epsins (Figure 3H). Importantly, we found elevated coronary artery disease (CAD) protective scores from Alb^hi^ hepatocytes towards Ldlr^hi^ hepatocytes that are positively associated with Ldlr expression (Figures 3G, 3H, 3I).

The livers from ApoE^-/-^/Liver-DKO mice exhibited elevated expression of other genes related to LDL/VLDL particle clearance including the VLDLr and Lrp1 (Figure 3C, 3D, Figure S9A). Importantly, the ApoE^-/-^/Liver-DKO mouse livers exhibited increased gene expression of proteins relevant to hepatic HDL cholesterol uptake including Scarb1 and LIPG^30^ (Figure 3C, 3D, Figure S9A). In addition, other genes operating in lipoprotein metabolism such as apoC3, apoC2, apoM, and LCAT were significantly downregulated in ApoE^-/-^/Liver-DKO hepatocytes (Figure 3C, 3E, Figure S9B).

We examined the relationship between Epsins1/2 expression and the protein levels of Ldlr in WT hepatocytes (Figures 4A-4D). Immunofluorescence and western blotting analyses of the effects of Epsins1/2 expression on Ldlr receptor levels in hepatocytes from Liver-DKO versus WT mice consuming a chow diet revealed modest increases in Ldlr protein (Figure 4A,4B). In contrast, the hepatocytes from Liver-DKO versus WT mice consuming a western diet had markedly increased Ldlr levels (Figures 4C,4D) that was likely due to the substantial upregulation of both Epsin1 and Epsin 2 in hepatocytes from WT mice consuming a western versus a chow diet (Figures 4E,4F). As apoE accelerates LDL/VLDL uptake via LDLr, VLDLr, Lrp1, and SR-BI^31,32^,we examined the effects of hepatic deletion of Epsins1/2 on apoE levels in WT mice and found that deletion of hepatic Epsins1/2 markedly increased hepatocyte apoE protein levels (Figures 4G,4H). In addition, both single cell and real-time quantitative PCR analyses revealed that deletion of Epsins1/2 remarkably increased expression of the apoAIV gene in hepatocytes from ApoE^-/-^/Liver-DKO versus ApoE^-/-^ mice (Figures 4I), which is highly relevant as studies have shown that apoAIV facilitates LDL clearance as well as reduces liver fat and gluconeogenesis ^33–35^. Indeed, deletion of hepatic Epsins1/2 increased the expression of apoAIV in all Alb^hi^ hepatocyte populations (Figure S9C). Studies with hepatocytes from Liver-DKO versus WT mice confirmed a marked increase in apoAIV protein (Figures 4J, 4K).

**Figure 4:**
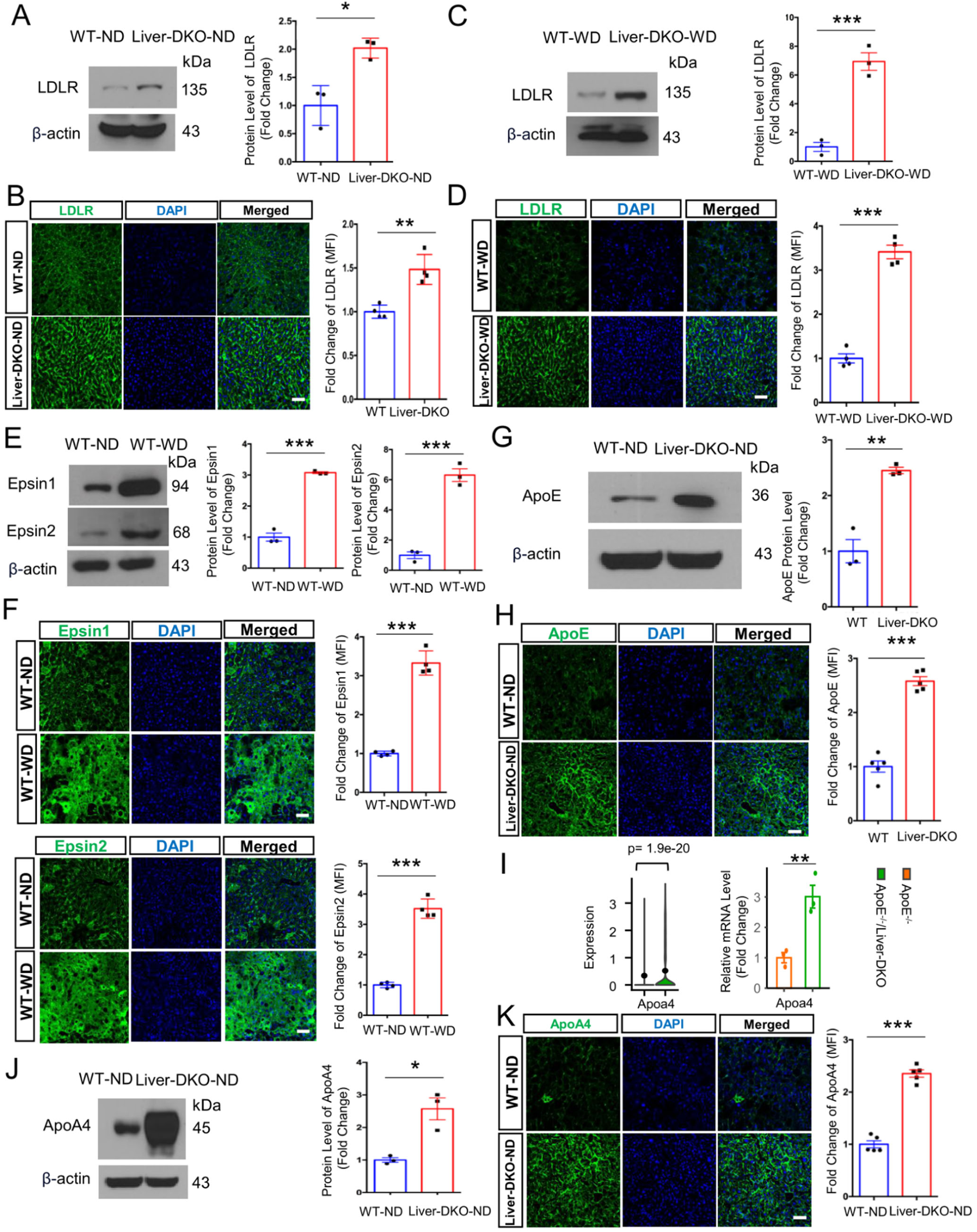
Elevated LDLR, ApoE and ApoA4 expression in the liver of liver-specific epsins deficiency mice and elevated Epsin1 and Epsin2 in the liver of WD-fed mice. A: Western blot (WB) analysis of liver tissue harvested from WT and Liver-DKO mice (normal diet, ND) revealed elevated LDLR expression in Liver-DKO mice (left). Data quantification of LDLR expression (right) (n=3, ***p< 0.001). B: Immunofluorescence (IF) analysis of LDLR expression level in liver cryosections from ND-fed WT, Liver-DKO mice (left), demonstrating elevated LDLR protein level in the livers of Liver-DKO. Mean fluorescence intensity (MFI) quantification of LDLR under ND (right) (n=4, ***p<0.001). Scale bar, 50 μm. C: Western blot (WB) analysis of liver tissue harvested from WT and Liver-DKO (western diet, WD) mice showed elevated LDLR expression in WD-fed Liver-DKO mice (left). Data quantification of LDLR expression (right) (n=3, ***p<0.001). D: Immunofluorescence (IF) analysis of LDLR expression in liver cryosections from WD-fed WT and Liver-DKO mice (left), demonstrating that elevated LDLR protein level in WD-fed Liver-DKO mice. Mean fluorescence intensity (MFI) quantification of LDLR under WD (right) (n=4, ***p<0.001). Scale bar,50 μm. E: Western blot (WB) analysis of liver tissue harvested from WT (normal diet, ND) and WT (western diet, WD) mice showed elevated epsin1 and epsin2 expression in WD-fed WT mice (left). Data quantification of Epsin1 and Epsin2 expression (right) (n=3, ***p<0.001). F: Immunofluorescence (IF) analysis of Epsin1 and Epsin2 expression in liver cryosections from ND-fed WT and WD-fed WT mice (left), demonstrating that both Epsin1 and Epsin2 are significantly induced in WD-fed WT mice. Mean fluorescence intensity (MFI) quantification of Epsin1 and Epsin2 under WD (right) (n=4, ***p<0.001). Scale bar,50 μm. G: Western blot analysis for the evaluation of ApoE expression in the livers in both ND-fed WT and Liver-DKO mice (left). Data quantification of ApoE expression (right) (n=3, ** p < 0.01). H: Immunofluorescence (IF) analysis of ApoE expression in liver cryosections from both ND-fed WT and Liver-DKO mice (left). Mean fluorescence intensity (MFI) quantification of ApoE between WT and Liver-DKO mice (right) (n=5, *** p < 0.001). Scale bar, 50 μm. I: Violin plot of Apoa4 by single cell analysis (left) and RT-qPCR evaluation of Apoa4 mRNA expression between ApoE^-/-^ controls and ApoE^-/-^ /Liver-DKO mice (n=3, **p<0.01). J: Western blot of ApoA4 for the liver lysates from ND-fed WT and Liver-DKO, β-actin is used as an internal reference (left), quantification data of ApoA4 protein level (right) (n=3, *p<0.05). K: Immunofluorescence (IF) analysis of ApoA4 in the livers of both ND-fed WT and Liver-DKO mice (left). Quantification of mean fluorescence intensity (MFI) of ApoA4 protein level (right). Scale bar, 50 μm. All statistical analysis (A-K) in WT-ND and Liver-DKO-ND comparison or WT-WD and Liver-DKO-WD comparison or WT-ND and WT-WD comparison is conducted by the Student’s t-test.

### Decreased Hepatic Lipogenesis and Strengthened Glycogen Biosynthesis in the Livers of Hepatic Epsin1/2 Deficient Mice

A closer examination by single cell or real-time quantitative PCR analysis of the effects of hepatic Epsins1/2 deletion on lipogenesis in ApoE^-/-^ mice demonstrates that the expression of genes known to suppress lipogenesis including Hnf4α, Sdc4, Rorα, LIPA, and PPARα (Figures 5A, Figures S9A, S10A, S10C) were markedly increased in livers of ApoE^-/-^/Liver-DKO versus ApoE^-/-^ mice. Correspondingly, detailed analyses revealed upregulated *Ror*α-(i.e., DHCR24) and *Sdc4*-(Fn1) associated anti-lipogenic signaling pathways in ApoE^-/-^ /Liver-DKO mice (Figure S10A)^36^. Importantly, ApoE^-/-^/Liver-DKO versus ApoE^-/-^ mouse livers exhibited diminished mRNA levels of genes that enhance hepatic lipogenesis including *Acaca*, *Scd1*, *Acly*, *Hmgcr,* and *Fasn* (Figures 5B-5C). In addition, the *Scd1* and *Acaca* mRNA levels decreased across all hepatocyte subpopulations (Figure S9C, Figure 5B). Furthermore, there was corresponding downregulation of prolipogenic Ppia-Bsg signaling pathway (Figures S10B). Importantly, western blotting and immunofluorescence analyses demonstrate that the levels of the fatty acid and cholesterol synthetic enzymes, Fasn and Hmgcr, were decreased in the livers of Liver-DKO versus WT mice (Figures 5D-5F). It is also worth noting that some genes linked to the suppression of lipogenesis such as *Nr1h4* and apoM were downregulated in ApoE^-/-^/Liver-DKO versus ApoE^-/-^ mouse livers ^37^ (Figure S9B, S10B). However, the overall effect of hepatic Epsins1/2 deletion was to markedly reduce liver fat as demonstrated by decreased Oil-Red-O staining of livers from ApoE^-/-^/Liver-DKO versus ApoE^-/-^ mice fed a western diet (Figure 5G).

**Figure 5:**
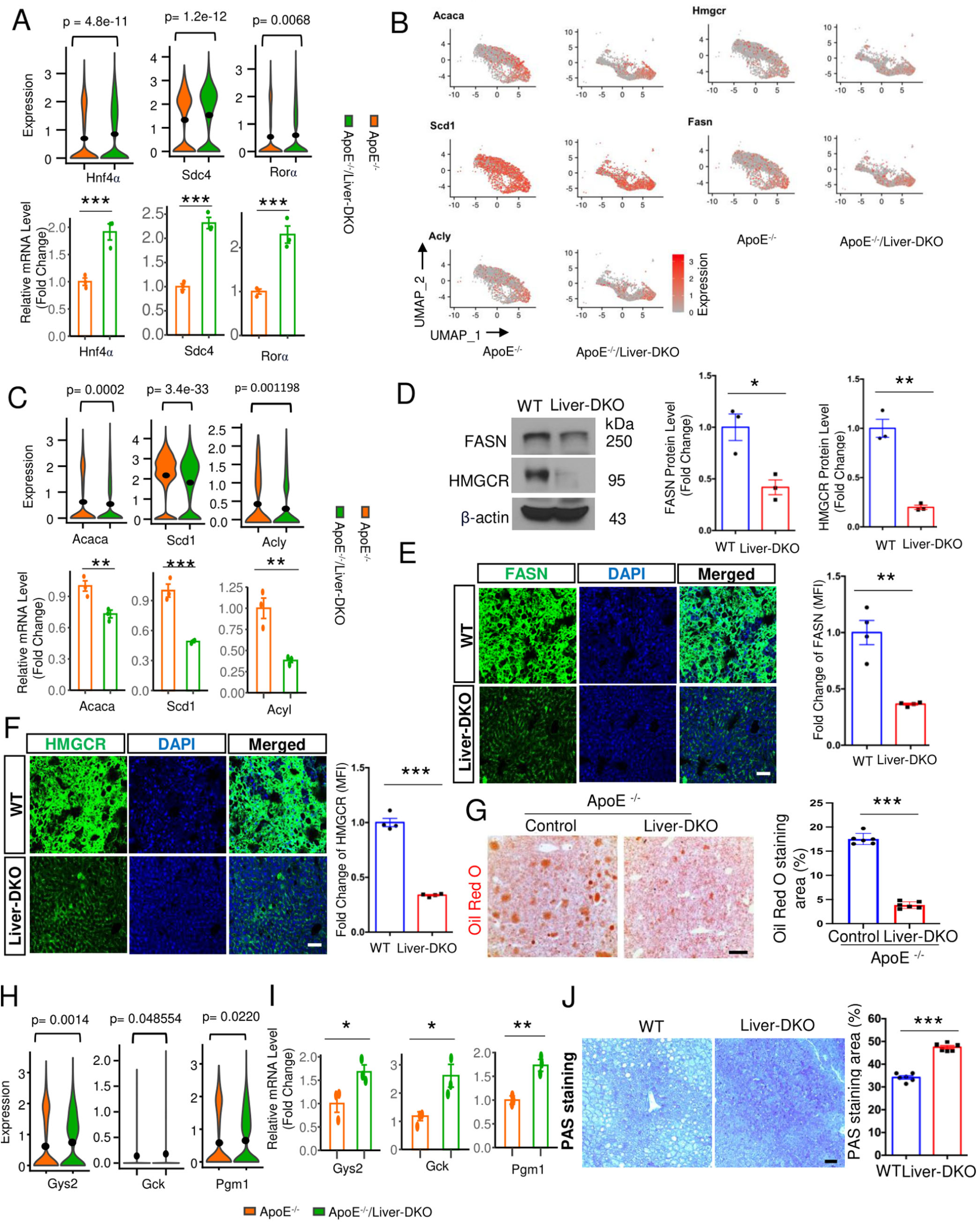
Diminished lipogenic gene expression in the liver of liver-specific epsins deficiency mice resulted in lower hepatic lipid accumulation and elevated glycogenic gene expression in the liver of Liver-DKO mice that induced higher hepatic glycogen. A: Violin plots revealed elevated expression of genes that suppression of lipogenesis (Hnf4α, Sdc4, Rorα) in ApoE^-/-^/Liver-DKO as compared to ApoE^-/-^, shown through single-cell analysis (left). Validation of Hnf4α, Sdc4, Rorα gene expression involved in inhibition of lipogenesis by real-time quantitative PCR (qPCR) in the livers from both ApoE^-/-^ and ApoE^-/-^/Liver-DKO mice (right) (n=3, *** p<0.001). B: Feature plots show diminished expression of lipogenic genes in ApoE^-/-^/Liver-DKO mice compared with ApoE^-/-^ mice, such as *Acaca*, *Scd1*, *Acly*, *Hmgcr*, *Fasn*. C: Violin plots of gene expression related to lipogenesis (*Acaca*, *Scd1*, *Acly*) in ApoE^-/-^/Liver-DKO as compared to ApoE^-/-^, shown through single-cell analysis (left). Validation of *Acaca*, *Scd1* and *Acly* gene expression involved in lipogenesis by real-time quantitative PCR (qPCR) in the livers from both ApoE^-/-^ and ApoE^-/-^/Liver-DKO mice (right) (n=3, **p<0.01, *** p<0.001). D: Western blotting analysis of FASN and HMGCR proteins in the liver lysates from WT and Liver-DKO mice (left), and the relative expression of FASN and HMGCR proteins in the liver of WT and Liver-DKO mice are quantified (right) (n=3, * p<0.05, **p<0.01). Diminished FASN and HMGCR protein levels in the liver lysates in Liver-DKO mice. E-F: Immunofluorescence analysis of FASN (E) and HMGCR (F) expression in the livers of both WT and Liver-DKO mice (left). MFIs of FASN and HMGCR are quantified (right) (n=4, **p<0.01, *** p<0.001). Diminished expression of FASN and HMGCR in the liver cryosections in Liver-DKO mice. Scale bar, 50 μm. G: ORO staining of the livers (left) from ApoE^-/-^ or ApoE^-/-^ / Liver-DKO mice fed a WD, and unpaired t-test (right) for lipid-stained areas (n=5, ***p< 0.001). Scale bar, 500 μm. H: Violin plots of gene expression related to glycogenesis (*Gys2*, *Gck*, *Pgm1*) in ApoE^-/-^/Liver-DKO as compared to ApoE^-/-^, shown through single-cell analysis. I: Validation of *Gys2*, *Gck* and *Pgm1*gene expression involved in glycogenesis by real-time quantitative PCR (qPCR) in the liver from both ApoE^-/-^ and ApoE^-/-^/Liver-DKO mice (n=3, * p <0.05, ** p<0.01). J: Periodic acid-Schiff’s (PAS) staining of the livers (left) from WT and Liver-DKO mice, and unpaired t-test (right) for glycogen-stained areas (n=6, ***p<0.001). Scale bar, 200 μm. Statistical analysis (A and C-J) in WT and Liver-DKO comparison or ApoE^-/-^ controls and ApoE^-/-^/Liver-DKO mice comparison is conducted by the Student’s t-test.

Cell-cell communication analysis using the MEBOCOST algorithm^25^ showed enhanced communication mediated by the cholesterol/Ldlr pathway in Ldlr ^hi^ hepatocytes in ApoE^-/-^/Liver-DKO mice (Figure S10C, S10D) and resulted in lowered mean abundance of cholesterol in ApoE^-/-^/Liver-DKO mice (Figure S10E). Interestingly, single cell and real-time quantitative PCR analyses also revealed enhanced expression of key genes that facilitate hepatic glycogenesis including Gck, Gys2, and Pgm1 in livers of ApoE^-/-^/Liver-DKO versus ApoE^-/-^ mice (Figures 5H, 5I). Intriguingly, the expression of glycogenesis genes (Pgm1 and Ugp2) was relatively more abundant in Ldlr ^hi^ and HC3 Alb^hi^ versus HC1 Alb^hi^ and HC2 Alb^hi^ hepatocyte (Figure S9C). Importantly, we observed a strengthened abundance of uridine diphosphate glucose (UDPG) as an intermediate metabolite for glycogenesis in Ldlr ^hi^ hepatocytes in ApoE^-/-^/Liver-DKO mice (Figure S10E), which is consistent with elevated glycogenesis gene expression in the livers of ApoE^-/-^/Liver-DKO mice. Indeed, Periodic acid-Schiff’s staining of livers in WT and Liver-DKO mice revealed a 39% increase in mean glycogen-stained areas in Liver-DKO mice compared with WT controls (Figure 5J). Overall, our data suggest that Alb^hi^ hepatocytes are prone to lipogenesis in the liver, while Ldlr^hi^ hepatocytes prefer glycogenesis, and that increased Epsins1/2 expression markedly impacts these processes in part by decreasing the propensity for hepatocyte conversion to Ldlr^hi^ cells.

### LDLR is Resistant to PCSK9-induced Proteasomal Degradation, Resulting in Elevated LDL Uptake in Liver-DKO mice, and LDLR Directly Binds to Epsin1 UIM Domain

We show that deletion of hepatic Epsins dramatically increases Ldlr protein levels (Figures 4A-4D) in comparison to Ldlr gene expression (Figures 3F-3G) suggesting that the Epsins impact Ldlr posttranslationally. As epsins are ubiquitin-binding endocytic adaptors, we hypothesized that epsins are required for PCSK9-mediated LDLR degradation. Indeed, western blot (WB) analyses of liver tissue harvested from WT and Liver-DKO mice injected with PCSK9-AAV8 (2x10^11^ genomes) revealed that the LDLR was markedly degraded by PCSK9 administration in WT mice, but not in the livers of Liver-DKO mice, with 3.2-fold more hepatic LDLR protein levels in Liver-DKO mice compared with WT controls (Figure 6A). In addition to liver tissue lysate analysis, we also performed WB analysis of lysates from primary hepatocytes isolated from WT and Liver-DKO mice, treated with PCSK9, cycloheximide (CHX) or MG132. We found PCSK9-induced LDLR degradation occurred independent of new protein synthesis (in the presence of CHX) but was blocked by either loss of epsins or the proteasomal inhibitor MG132 (Figure 6B). The protein levels of LDLR were quantified, with significantly higher LDLR expression in the hepatocytes from Liver-DKO mice either with or without MG132 treatment (Figure 6B).

**Figure 6:**
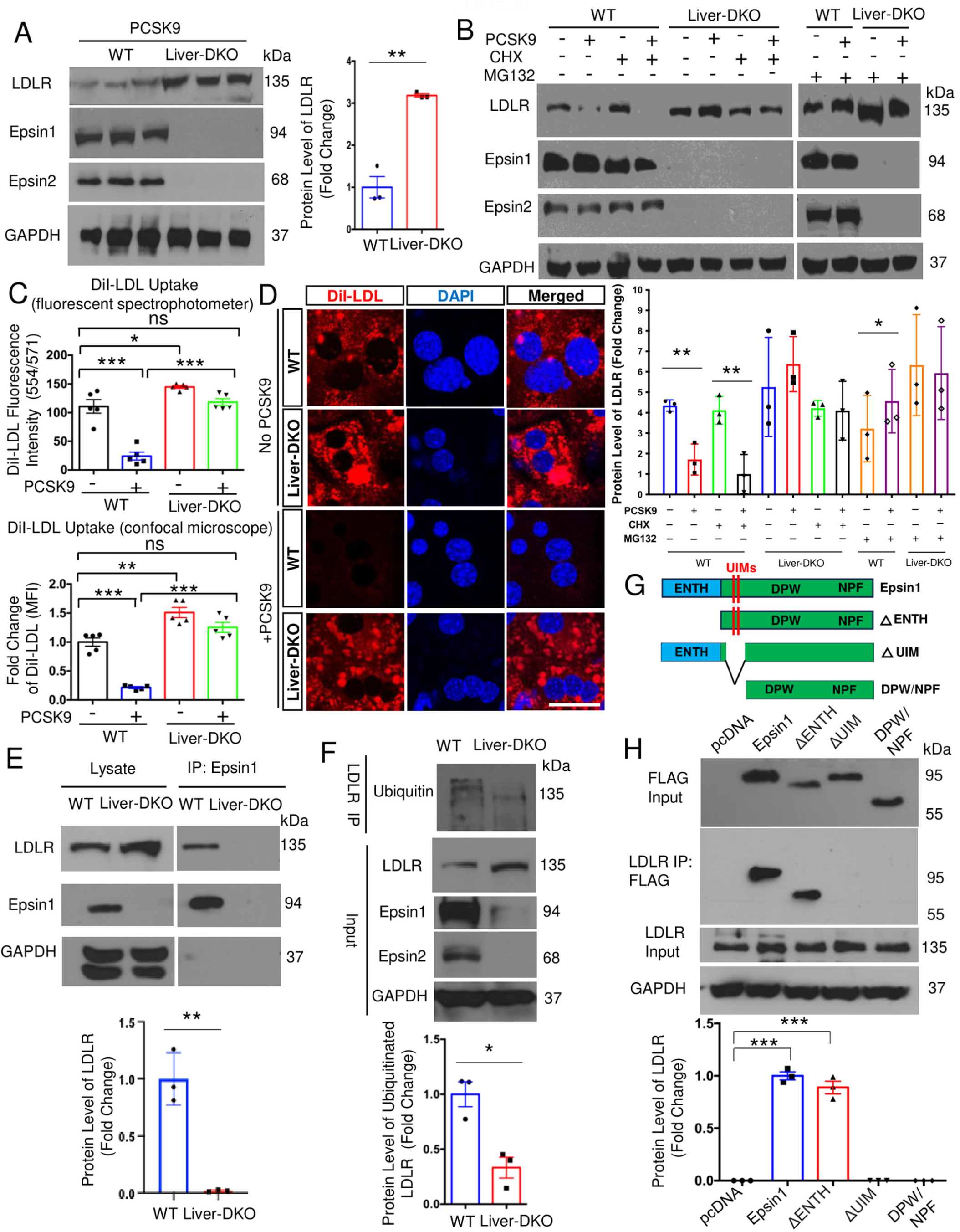
LDLR is Resistant to PCSK9-induced Proteasomal Degradation Results in Elevated LDL Uptake in Liver-DKO mice and LDLR Directly Binds to Epsin1 UIM Domain. A: Western blot (WB) analysis of liver tissue harvested from WT and Liver-DKO mice injected with PCSK9-AAV8 virus revealed PCSK9-triggered LDLR degradation is inhibited in Liver-DKO mice (left). Data quantification (right) (n=3, **p<0.01). B: WB analysis of lysates from primary hepatocytes isolated from WT and Liver-DKO mice, treated with PCSK9, cycloheximide (CHX) and MG132 showed PCSK9-induced LDLR degradation occurred independent of new protein synthesis (in the presence of CHX) but was blocked by either loss of epsins or proteasomal inhibitor MG132 (top). Data quantification (bottom) (n=3, **p<0.01 vs lane 1, *p<0.05 vs lane 2). C: DiI-LDL uptake assay for the hepatocytes isolated from WT and Liver-DKO mice with or without PCSK9 treatment by measuring the DiI-LDL fluorescence intensity using a fluorescent spectrophotometer (ex 554/ em 571). Diminished mean DiI-LDL fluorescence intensity is detected in WT hepatocytes under PCSK9 treatment, while PCSK9-induced hepatocytes from Liver-DKO mice maintain high levels of mean DiI-LDL fluorescence intensity (n=5, * p < 0.05, *** p < 0.001). D: DiI-LDL uptake assay for the hepatocytes isolated from WT and Liver-DKO mice with or without PCSK9 treatment by taking DiI-LDL fluorescence images using confocal microscope. The mean fluorescence intensity (MFI) of different groups was quantified (lower-left corner). PCSK9-treated WT hepatocytes exhibited significantly diminished MFI of DiI-LDL compared with WT hepatocyte controls, while PCSK9-treated Liver-DKO hepatocytes maintained high levels of MFI of DiI-LDL compared with WT hepatocyte controls (n=5, ** p < 0.01, *** p < 0.001). Scale bar, 50 μm. E: Epsin1 antibody immunoprecipitation with lysates from liver tissue in both WT and Liver-DKO mice, anti-LDLR antibody was used to detect the LDLR expression between WT and Liver-DKO mice after IP assay. Liver lysates from both WT and Liver-DKO mice were also used as input control for testing anti-LDLR, Epsin1, and GAPDH antibodies (top). Data quantification of IP assay (bottom) (n=3, **p<0.01). Anti-epsin1 co-IPs showed LDLR directly binds Epsin 1 in WT, but not in Liver-DKO mice. F: LDLR antibody immunoprecipitation with lysates from liver tissue in both WT and Liver-DKO mice, anti-Ubiquitin antibody was used to detect the ubiquitinated LDLR between WT and Liver-DKO mice. Liver lysates from both WT and Liver-DKO mice were also used as input control for testing anti-LDLR, Epsin1, Epsin2, and GAPDH antibodies (top). Data quantification of IP assay (bottom) (n=3, *p<0.05). G: Schematic of Epsin 1 deletion mutants and individual protein domains, including full-length epsin1, ΔENTH, ΔUIM, DPW/NPF plasmids. H: LDLR antibody immunoprecipitation with lysates from HepG2 cells that transfected with FLAG-fused plasmids mentioned above. An anti-FLAG antibody was used to detect FLAG expression. HepG2 cell lysates that transfected with different FLAG-fused plasmids were also used as input control for testing anti-FLAG, LDLR, and GAPDH antibodies (top). Data quantification of IP assay (bottom) (n=3, ***p<0.001). Statistical analysis (A, E, F, H) is conducted by the Student’s t-test, and statistical analysis (B, C, D) is conducted by two-way ANOVA with post hoc test.

Next, we examined the effects of Epsin1/2 expression on the uptake of DiI-LDL by fluorescence spectrophotometer (Figure 6C) or by confocal microscopy analyses (Figure 6D). There was a 31% increase in mean DiI-LDL uptake in Liver-DKO hepatocytes compared with WT controls (Figure 6C). Consistent with the PCSK9-AAV8-induced LDLR degradation, a 78% decrease DiI-LDL uptake was observed in PCSK9-AAV8-infected WT control hepatocytes versus WT controls (Figure 6C) by spectrophotometer analysis. Moreover, the analysis revealed a 4.9-fold more uptake of DiI-LDL in PCSK9-AAV8-infected Liver-DKO hepatocytes compared with PCSK9-AAV8-infected WT controls (Figure 6C). In addition, confocal microscopy analysis revealed a 51% increase in uptake of DiI-LDL in the hepatocytes in Liver-DKO mice compared with WT controls (Figure 6D). Importantly, confocal microscopy analysis demonstrates that PCSK9-AAV8-infected Liver-DKO hepatocytes had 5.8-fold more uptake of DiI-LDL compared with PCSK9-AAV8-infected WT controls (Figure 6D). Taken together, the hepatocytes in Liver-DKO mice possessed elevated capacity for LDL-C clearance compared with hepatocytes in WT controls.

To study the interaction between LDLR and Epsin1, we performed an anti-epsin1 co-immunoprecipitation (co-IPs) analysis. We found LDLR directly binds epsin1 in WT, but not in Liver-DKO primary hepatocytes (Figure 6E). In addition, we discovered that levels of ubiquitinated LDLR were significantly diminished in the liver lysate from Liver-DKO mice by testing ubiquitin expression after LDLR antibody for immunoprecipitation (IP) (Figure 6F). To study which epsin1 motif can bind to LDLR, we transfected different FLAG-tagged epsin1 deletion mutants’ plasmids into HepG2 cells (Figure 6G), including pcDNA (empty plasmid), full-length epsin1 plasmid, epsin1-ΔENTH plasmid, epsin1-ΔUIM plasmid, epsin1-DPW/NPF plasmid. Intriguingly, we found that LDLR can specifically bind to both the full-length epsin1 and epsin1-ΔENTH but does not bind to epsin1-ΔUIM and epsin1-DPW/NPF, indicating the epsin1-UIM domain is the binding motif for the interaction between epsin1 and LDLR (Figure 6H).

### Inhibited Low-density Lipoprotein Particle Clearance, High-density Lipoprotein Assembly, and Glycogen Biosynthesis in the Liver of PCSK9-D374Y Mutant Mice

Overexpression of PCSK9 induces hyperlipidemia and accelerates atherosclerotic plaque formation^13^. We show that deletion of hepatic Epsins1/2 negates the effects of PCSK9 overexpression on atherosclerosis (Figure 6). In addition, deletion of Epsins 1/2 negates hepatic PCSK9 mediated proteasomal degradation of the LDLr. Therefore, we further explored the underlying hepatic signaling pathways in relation to PCSK9 and Epsins by analyzing liver single-cell RNA-seq data from PCSK9-D374Y mutation mice. PCSK9-D374Y gain-of-function mutant has a markedly increased affinity for LDLR and promotes its degradation^38^. UMAP visualization showed two distinct hepatocyte subclusters between control and PCSK9-D374Y mutant (Figure S11A), namely, HC Alb^hi^ hepatocytes and HC Ldlr^hi^ hepatocytes, based on the expression of hepatocyte markers (Figure S12A). As expected, we identified diminished Ldlr expression in all hepatocytes in PCSK9-D374Y mutant (Figure S11B, S11K). Furthermore, we discovered elevated epsin1 expression in all hepatocytes in PCSK9-D374Y mutant (Figure S11C). Subsequent analyses revealed the downregulated pathways of low-density lipoprotein particle clearance and the upregulated pathways of the glycolytic process in PCSK9-D374Y mutant (Figure S11D-S11F, S13A, S13B). Moreover, we found elevated expression of lipogenic genes, e.g., *Acly*, *Fasn*, and diminished expression of glycogenic genes, e.g., *Gys2*, *Ugp2*, in PCSK9-D374Y mutant (Figure S12E). Importantly, almost all the apolipoprotein genes and genes that control HDL assembly (*Abca1* and *Apoa1*), cholesterol efflux (*Apoa1*, *Apoa2* and *Apoa4*) and the final step of reverse cholesterol transport (*Abcg5*, *Abcg8*) are dramatically inhibited in PCSK9-D374Y mutant (Figure S12C, Figure S12D), implying diminished HDL assembly and curtailed cholesterol clearance.

Further RNA velocity analysis demonstrated an inhibited propensity for hepatocyte differentiation from HC Alb^hi^ hepatocytes towards HC Ldlr^hi^ hepatocytes in the presence of elevated hepatic epsins in PCSK9-D374Y mutant (Figure S11G, Figure S12B). Consequently, we discovered lowered CAD protective scores in both HC Alb^hi^ and HC Ldlr^hi^ hepatocytes in PCSK9-D374Y mutants (Figure S11H). Metabolite analyses using the MEBOCOST algorithm showed weakened communication interactions for cholesterol/Ldlr pathway in all hepatocytes in PCSK9-D374Y mutant (Figure S11I). Subsequently, we found a lowered mean abundance of uridine diphosphate glucose (UDPG) as an intermediate metabolite for glycogenesis in all hepatocytes in PCSK9-D374Y mutant (Figure S11J).

### Nanoparticle-mediated Delivery of Epsins siRNAs Potently Inhibits Lesion Development, Reduces Foam Cell Formation, and Decreases Cholesterol and TG Levels in ApoE^-/-^ Mice

To explore the therapeutic potential of targeting hepatic epsins, we employed nanoparticle-encapsulated siRNAs specifically targeting epsins in the liver. Therapeutically, we applied nanoparticles (NPs) encapsulated epsin1 and epsin2 siRNAs (Figure 7A) to inject ApoE ^-/-^ mice under WD-treatment. Control siRNA NPs were injected into WD-fed ApoE ^-/-^ mice as the control. Western blots of liver lysates isolated from WD-fed ApoE ^-/-^ mice (8 weeks) that were treated with control or epsins siRNA NPs revealed dramatically diminished epsin1 and epsin2 protein expression, indicating the high efficiency of epsins siRNA NPs for knockdown of epsin1 and epsin2 proteins (Figure 7B). Mean serum cholesterol and triglyceride (TG) levels showed striking 31% and 22% decreases in epsin1 and epsin2 siRNAs NPs treated WD-fed ApoE ^-/-^ mice compared with control siRNA NPs treated WD-fed ApoE ^-/-^ mice (Figure 7C,7D). Importantly, epsin1 and epsin2 siRNAs NPs treated WD-fed ApoE ^-/-^ mice exhibited a 60% reduction in *en face* aortic lesion areas and a 62% reduction in lesions of the aortic roots compared with control siRNA NPs treated WD-fed ApoE ^-/-^ mice (Figure 7E, 7F). In addition, we discovered remarkably diminished mean fluorescence intensity (MFI) of CD68 in epsin1 and epsin2 siRNAs NPs treated WD-fed ApoE ^-/-^ mice compared with control siRNA NPs treated WD-fed ApoE ^-/-^ mice, indicating lowered macrophage accumulation in the atherosclerotic lesions in epsin1 and epsin2 siRNAs NPs treated ApoE ^-/-^ mice (Figure 7G). These findings suggest that targeting liver epsins presents a novel and promising therapeutic strategy for the treatment of atherosclerosis.

**Figure 7:**
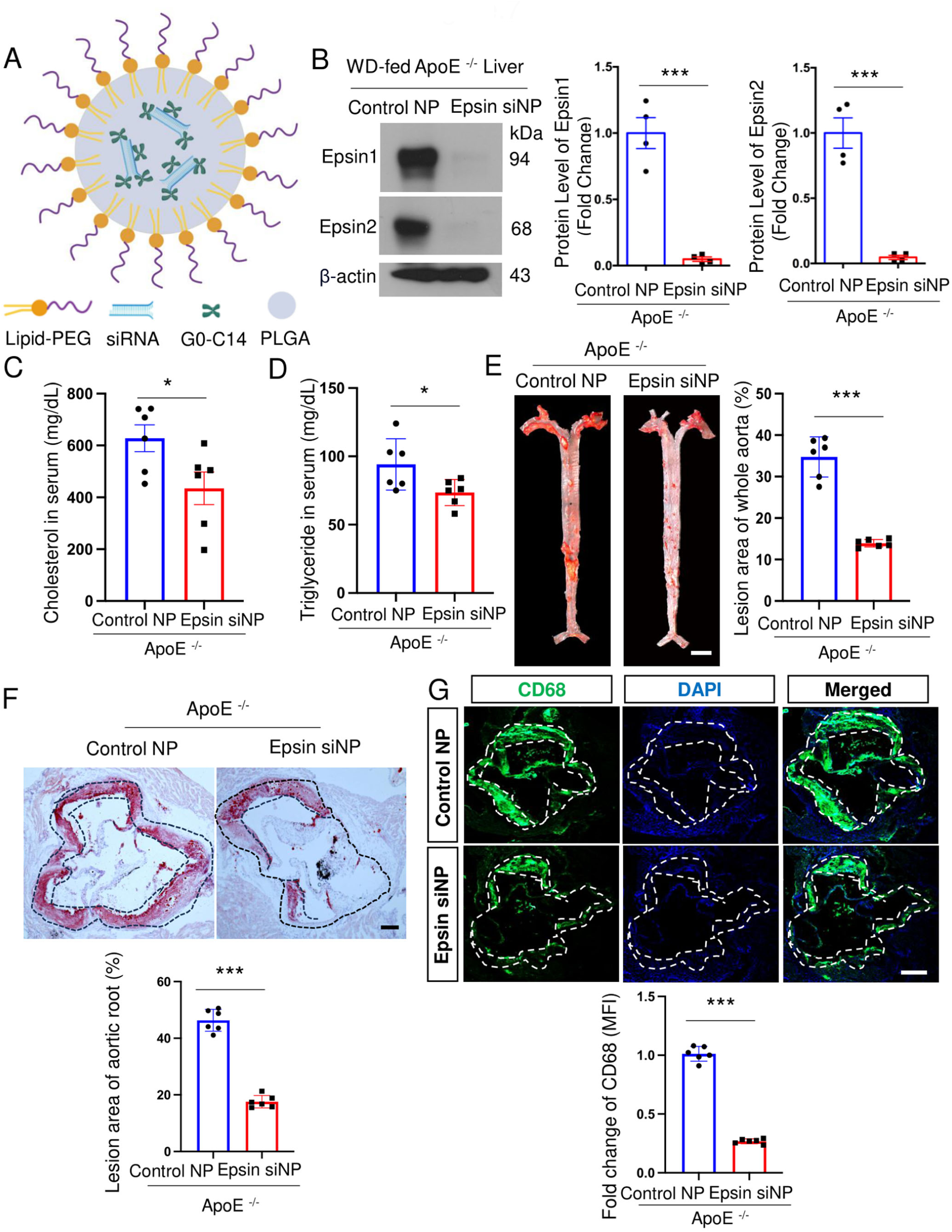
Nanoparticles (NPs) with epsin1/2 siRNA inhibit lesion formation and macrophage accumulation. A: Schematic of the siRNA encapsulated lipid nanoparticles (LNP) platform composed of a lipid-PEG shell with a PLGA core containing G0-C14/siRNA complexes. B: Western blots of liver lysates were isolated from WD-fed ApoE^-/-^ mice (8 weeks) and treated with control or epsin siRNA NPs (1 nmoles) for 3 weeks (2 doses/week) (left). The protein expression of Epsin1 and Epsin2 under control or epsin siRNA NPs treatment was quantified (right) (n=4, *** p<0.001). C-D: Plasma cholesterol (C) and triglyceride (TG) levels (D) in WD-fed ApoE^-/-^ (WT) and epsin siRNA-nanoparticle (NP)-treated mice (n=6, *p<0.05). E: *En face* ORO staining of aortas (left) from control siRNA NP-treated ApoE^-/-^ or epsin1/2 siRNA NP-treated ApoE^-/-^ mice fed a WD, and unpaired t-test (right) for lesion areas (n=6, ***p<0.001). Scale bar, 5mm. F: Aortic roots from control siRNA NP-treated ApoE^-/-^ or epsin1/2 siRNA NP-treated ApoE^-/-^ mice were stained with ORO (top), and quantification of aortic root lesion areas (bottom) (n=6, ***p<0.001). Scale bar, 200 μm. G: Aortic roots from control siRNA NP-treated ApoE^-/-^ or epsin1/2 siRNA NP-treated ApoE^-/-^ mice were stained by macrophage marker, CD68 antibody (top), and the mean fluorescence intensity (MFI) of CD68 is quantified (bottom) (n=6, ***p<0.001). Scale bar, 300 μm. Statistical analysis (B-G) in Control NP and Epsin siNP comparison is conducted by the Student’s t-test.

## DISCUSSION

In this study, we found that epsin1 and epsin2 expression dramatically increased by 98% and 68%, respectively, in the aortas in ASCVD patients compared with healthy controls (Figure S3A-S3C), suggesting the expression of epsin1 and epsin2 is positively correlated with atherosclerosis. Our previous studies have elucidated an atheroprone function of epsins in endothelial cells^4^, macrophages^16,17^, and vascular smooth muscle cells^39^ due to significantly elevated inflammation in the atherosclerotic plaque. However, atherosclerosis is initiated by the abnormal accumulation of LDL-C in the subendothelial layer of the arterial wall due to hyperlipidemia^40^. The liver is the central organ for the control of cholesterol homeostasis^41^ and controls blood levels of LDL-C by regulating LDLR expression, which mediates hepatic clearance of circulating LDL-C. Lowering LDL-C remains the most effective approach for prevention of ASCVD^12^. PCSK9 is dominantly expressed in the liver, and PCSK9 binds to LDLR and targets the LDLR for degradation^42^. Monoclonal antibody and interference-RNA therapies that target PCSK9 are extremely effective in reducing circulating levels of LDL-C ^9,10^; however, the mechanistic details of PCSK9-mediated LDLR degradation remain insufficiently understood.

In this study, our scRNA-seq data analyses in the context of aortas revealed 22% decrease in cell proportion of macrophages in the aortas in ApoE^-/-^/Liver-DKO mice strongly supports that attenuated inflammation ameliorates atherosclerosis^43^ (Figure S4C, Figure 2A). Furthermore, a 45% decrease in cell proportion of fibroblasts in the aortas in ApoE^-/-^/Liver-DKO mice as compelling evidence of reduced collagen accumulation that is consistent with the dramatic reduction in atherosclerotic lesion size^44^ (Figure S4C, Figure 2D). In addition, a 91% increase in endothelial cell proportion in the aortas in ApoE^-/-^/Liver-DKO mice is consistent with the upregulation of endothelial cell differentiation and downregulation of endothelial cell activation and migration and downregulated pathways involved in inflammation in macrophages likely reduce the progression of atherosclerosis^45,46^ (Figure S4C, Figure S5A, Figure 1E-1G). Notably, a 63% increase in vascular smooth muscle cells (VSMCs) proportion in the aortas in ApoE^-/-^/Liver-DKO mice with substantially elevated expression of markers of the contractile VSMCs that inhibited phenotypic switching into synthetic VSMCs that impeded atherosclerosis^47^ (Figure S4C, S4D, Figure 2A, 2D). Hence, the markedly impeded atheroma in the aortas in ApoE^-/-^/Liver-DKO mice is partially attributed to increased cell proportion of healthy endothelial cells, reduced cell proportion of inflammatory macrophages, attenuated extracellular matrix and collagen production, and the enrichment of contractile VSMCs.

Hepatic loss of epsins resulted in lower plasma cholesterol and triglyceride levels in both WD-fed ApoE^-/-^/Liver-DKO mice and PCSK9-AAV8 injected Liver-DKO mice compared with their controls (Figure 1A, Figure S7A). Importantly, the plasma cholesterol and triglyceride levels are not impacted by epsins deficiency cell types, including endothelial cells, macrophages, and vascular smooth muscle cells^4,16,17,39^. Taken together, these findings indicate that epsins in the hepatocytes participate in lipoprotein metabolism. The liver is the main organ that regulates circulating LDL-C levels, in large part by regulating LDLR-mediated LDL-C clearance^41^. In this study, we identified higher LDLR expression in the livers of Liver-DKO mice than WT controls (Figure 4A-4D). Our single-cell RNA sequencing data showed upregulated pathway of low-density lipoprotein particle clearance and downregulated pathway of plasma lipoprotein particle assembly in the hepatocytes in ApoE^-/-^/Liver-DKO mice (Figure 3C-3E), which is consistent with their reduced plasma cholesterol and triglycerides. Subsequently, we exhibited evidence of enhanced communication score between LDLR and cholesterol in the hepatocytes in ApoE^-/-^/Liver-DKO mice by MEBOCOST analysis (Figure S10C-S10E), indicating boosted capacity of low-density lipoprotein cholesterol (LDL-C) clearance by LDLR in ApoE^-/-^/Liver-DKO mice. In addition, hepatic Epsins deficiency impacted lipogenesis, which is substantiated by markedly diminished expression of genes and proteins that are involved in lipogenesis in ApoE^-/-^/Liver-DKO mice (Figure 5A-5F). Furthermore, hepatocyte Epsins deficiency was associated with increased differentiation of Alb^hi^ hepatocytes towards Ldlr^hi^ hepatocytes in ApoE^-/-^/Liver-DKO mice compared with ApoE^-/-^ controls (Figure 3H), resulting in a 59% increase in the proportion of Ldlr^hi^ hepatocytes (Figure 3B). Consequently, we found a dramatically reduced atherosclerotic lesion area in ApoE^-/-^/Liver-DKO mice compared with ApoE^-/-^ controls (Figure 1B, 1C). In addition to enhanced LDL-C clearance, we found the elevated UDPG metabolite level further promotes hepatic glycogenesis in Ldlr^hi^ hepatocytes in ApoE^-/-^/Liver-DKO mice (Figure S10E). Chen *et al.* recently reported that hepatic glycogenesis inhibits lipogenesis^48^. Accordingly, the enhanced glycogen level in the liver inhibits fatty acid synthesis that could also contribute to reduced cholesterol and triglyceride in plasma in ApoE^-/-^/Liver-DKO mice.

Currently, no single-cell transcriptomics study of human livers from patients in the context of coronary artery disease is available, and most single-cell RNA sequencing studies for patients in the context of coronary artery disease are for exploring the transcriptome differences of atherosclerotic plaques^49,50^. Therefore, it is impossible to compare our liver single-cell transcriptome data with counterparts from coronary artery disease patients. However, a recently published article explored the liver transcriptome by single-cell RNA sequencing for PCSK9-D374Y mice with an emphasis on the analysis of Kupffer cells^51^; this dataset empowered us to analyze the heterogeneity of hepatocytes in the livers between PCSK9-D374Y mice and controls. The PCSK9-D374Y mutation potently elevates the affinity between PCSK9 and LDLR interaction, and further promotes the degradation of the LDLR^52^. In this study, we reanalyzed the liver single-cell transcriptome dataset by emphasizing the analysis on hepatocytes in PCSK9-D374Y mice^51^. Intriguingly, we found an inhibited propensity for differentiation of Alb^hi^ hepatocytes towards Ldlr^hi^ hepatocytes in PCSK9-D374Y mice compared with controls that cause weakened glycogenesis, enhanced lipogenesis, and lowered CAD protective score (Figure S11G, S11H, Figure S12E). Accordingly, there are a number of phenotypic similarities between WD-fed PCSK9-AAV8-injected control mice and WD-fed ApoE^-/-^ mice (Figure 1A-1C, Figure S7A-S7C). Mechanistically, we identified an elevated epsin1 expression level by 69% in Ldlr^hi^ hepatocytes that might cause impeded clearance of LDL particles and VLDL particles by curtailment of LDLR expression level by 72% in Ldlr^hi^ hepatocytes in PCSK9-D374Y mice (Figure S11B, S11C). Subsequently, MEBOCOST analysis revealed a diminished communication score between LDLR and cholesterol in the hepatocytes in PCSK9-D374Y mice (Figure S11I-S11K), which is compatible with serum cholesterol accumulation due to reduced hepatic LDLR levels in PCSK9-D374Y mice^53^. Similar to the phenotype in ApoE^-/-^ mice, decreased UDPG expression in hepatocytes in PCSK9-D374Y mice (Figure S11J) and curbed glycogenesis further contribute to elevated lipogenesis^48^. In summary, two different mouse models of atherosclerosis, ApoE^-/-^ and PCSK9-D374Y mice, shared similar hepatocyte heterogeneity and common pathological signaling pathways for inducing atherosclerosis or dyslipidemia, which might be highly correlated with epsins-mediated LDLR degradation in the liver.

We demonstrated how liver epsins regulate PCSK9-mediated LDLR degradation in the liver (Figure 8). In liver epsins-deficient mice, LDLR expression is markedly increased in the membrane of hepatocytes, promoting enhanced LDL-C clearance (Figure 8). PCSK9 protein mediates LDLR protein degradation, reducing hepatic uptake of LDL-C^54^. In this study, when PCSK9 is overexpressed by injection of PCSK9-AAV8 into WT controls and Liver-DKO mice, the LDLR is remarkably degraded in WT controls; however, the degradation of LDLR is significantly inhibited in Liver-DKO mice, with a 2.2-fold increase in LDLR expression level (Figure 6A), which demonstrates that liver epsins are essential for PCSK9- mediated LDLR degradation. In studies of primary hepatocytes isolated from WT and Liver-DKO mice were treated with PCSK9, cycloheximide (CHX) or proteasomal inhibitor MG132, we discovered that PCSK9-induced LDLR degradation occurred independent of new protein synthesis, and LDLR degradation was blocked by loss of epsins in the liver (Figure 6B). Importantly, 3.9- or 4.8-fold increase in the uptake of DiI-LDL in mean by the PCSK9-treated hepatocytes in Liver-DKO mice compared with PCSK9-treated WT controls is associated with the inhibition of PCSK9-mediated degradation of LDLR, which is consistent with elevated capacity for LDL particle clearance identified by single cell RNA-sequencing (Figure 6C, 6D). Subsequently, we discovered that epsin1 can directly bind to LDLR protein, which further triggers its degradation (Figure 6E). Our previous studies revealed that epsins are critical adaptor proteins that are involved in endocytosis^15,18,55^. To explore which motifs in epsin1 can specifically bind to LDLR, such as ENTH, UIM, DPW and NPF (Figure 6G), FLAG-tagged epsin1 deletion mutants’ plasmids have been transfected into HepG2 cells. Importantly, the LDLR binds specifically to the ubiquitin-interacting motif (UIM) motif but does not bind to other motifs (Figure 6H). The UIM in epsins facilitates ubiquitination-mediated protein degradation^56^. Consequently, diminished LDLR expression in the liver is due to the interaction between LDLR and the epsin1 UIM motif, which activates ubiquitination of LDLR, facilitating its degradation in WT controls, but not in Liver-DKO mice. In addition, the ubiquitinated LDLR was decreased in the liver lysate from Liver-DKO mice; consequently, LDLR expression was increased in the input lysate in Liver-DKO mice compared with WT controls (Figure 6F), demonstrating that the loss of epsins enhances the stability of LDLR. MG132 proteasome inhibitors blocked the degradation of LDLR by inhibition of ubiquitination^57^. In summary, epsins play gatekeeping roles in the PCSK9-mediated ubiquitination-driven LDLR degradation mediated by the interaction between epsin1 UIM motif and LDLR. Therefore, the liver epsins are potentially attractive targets for the treatment of dyslipidemia and atherosclerosis by protecting hepatic LDLRs from degradation.

**Figure 8:**
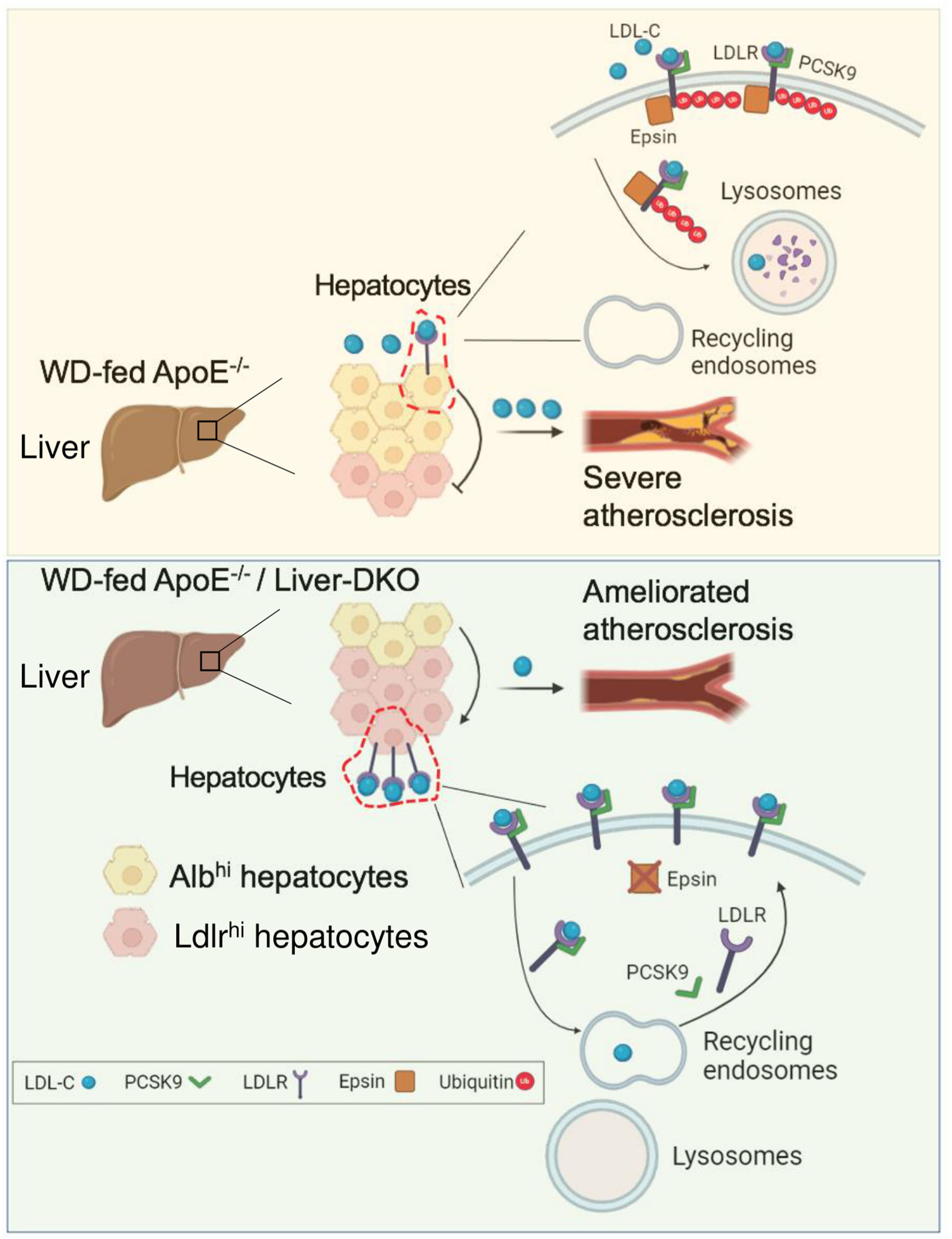
Western diet (WD)-fed ApoE^-/-^/Liver-DKO mice have elevated low-density lipoprotein cholesterol (LDL-C) clearance compared with WD-fed ApoE^-/-^ mice is attributable to their lower Alb^hi^ lipogenic hepatocytes and higher Ldlr ^hi^ glycogenic hepatocytes proportions. In the liver, glycogenesis inhibits lipogenesis. Consequently, the progression of atherosclerotic plaques is significantly ameliorated in WD-fed ApoE^-/-^ /Liver-DKO mice. Mechanistically, loss of epsins protein in the liver prevents ubiquitination-driven LDLR degradation. The expressed LDLR in hepatocyte cell membrane uptakes LDL-C from the circulation. In the presence of epsins protein in the liver (top), in WT mice, PCSK9 binds to LDLR, epsins protein mediates LDLR ubiquitination, and the ubiquitinated LDLR is directed to lysosomes for protein degradation. Consequently, elevated circulating LDL-C due to PCSK9-mediated LDLR degradation. In the absence of epsins protein in the liver (bottom), in epsin1/2 Liver-DKO mice, PCSK9 binds to LDLR, but the LDLR ubiquitination is abolished thanks to the deficiency of epsins protein. The LDLR is directed to recycling endosomes, and LDLR protein can be recycled and back to the plasma membrane of the hepatocytes to bind more LDL in the circulation and lower blood LDL-C levels.

The liver is the primary organ for lipid nanoparticle accumulation following intravenous administration^58^, and lipid nanoparticle-mediated RNAs or siRNAs delivery holds great potential for treating liver diseases^59,60^. In this study, targeting liver epsins with lipid nanoparticle encapsulated siRNAs was highly efficacious at inhibiting dyslipidemia and impeding atherosclerosis. We discovered notably 31% reduced cholesterol and a 22% reduction of triglyceride levels in the epsin1 and epsin2 siRNA NPs injected group compared with the control NPs group (Figure 7C, 7D). In addition, we found a 60% reduction in *en face* aortic lesion areas and a 62% reduction in lesions of aortic roots in epsin1 and epsin2 siRNA NPs-injected mice compared with the control NPs group (Figure 7E, 7F). These results support hepatic epsins as novel therapeutic targets for the treatment of hyperlipidemia and the prevention of atherosclerosis.

## Conclusions

Here, we discovered markedly diminished atherosclerotic lesions and reduced serum cholesterol levels in both WD-fed ApoE^-/-^/Liver-DKO mice and WD-fed PCSK9-AAV8 injected Liver-DKO mice. We demonstrated that the loss of epsins in the liver inhibits atherosclerosis by enhancing LDLR stability. Mechanistically, epsins bind LDLR via the ubiquitin-interacting motif (UIM), and PCSK9-triggered LDLR degradation was abolished by depletion of epsins, dramatically lowering plasma cholesterol levels and preventing atheroma progression. Additionally, in the liver, the increased propensity of hepatocyte differentiation from Alb^hi^ towards Ldlr^hi^ hepatocytes in the absence of hepatic epsins inhibits atheroma progression by enhancing LDL-C clearance. Targeting liver epsins with nanoparticle-encapsulated epsins siRNAs, was highly efficacious at reducing plasma lipid levels and impeding atherosclerosis. These findings provide us with a novel therapeutic strategy for the treatment of hyperlipidemia to combat atherosclerosis.

## Acknowledgments

We thank Dr. Jinjun Shi’s lab at Brigham and Women’s Hospital for the synthesis of epsin1 and espin2 siRNA nanoparticles. We thank the imaging core at Boston Children’s Hospital, and the Biopolymers Facility at Harvard Medical School for quality control analysis of DNA libraries prepared for scRNA-sequencing. We thank the animal core facility for the daily maintenance at Boston Children’s Hospital.

## Author contributions

B.Z., K.C., and H.C. conceived and designed the experiments. B.Z., K.C. performed most of the experiments. B.W. contributed to animal experiments. K.G. and K.C. analyzed the scRNA-seq data and performed bioinformatic work. M.M. and S.S. measured plasma cholesterol and triglycerides. S.W. performed the mouse genotyping and colony maintenance. B.Z., K.G., K.L., D.W., S.B., P.G.Y., D.B.C., M.F.L, W.W, C.A.B, K.C. and H.C. wrote and edited the article. All the authors reviewed and provided feedback on the article.

## Supplemental Material

WB Raw Films

Major Resources Table

Nonstandard Abbreviations and Acronyms

Clinical Perspective

Supplemental Figure 1-14

Table S1-S2

## Supplemental Figures

**Figure S1:**
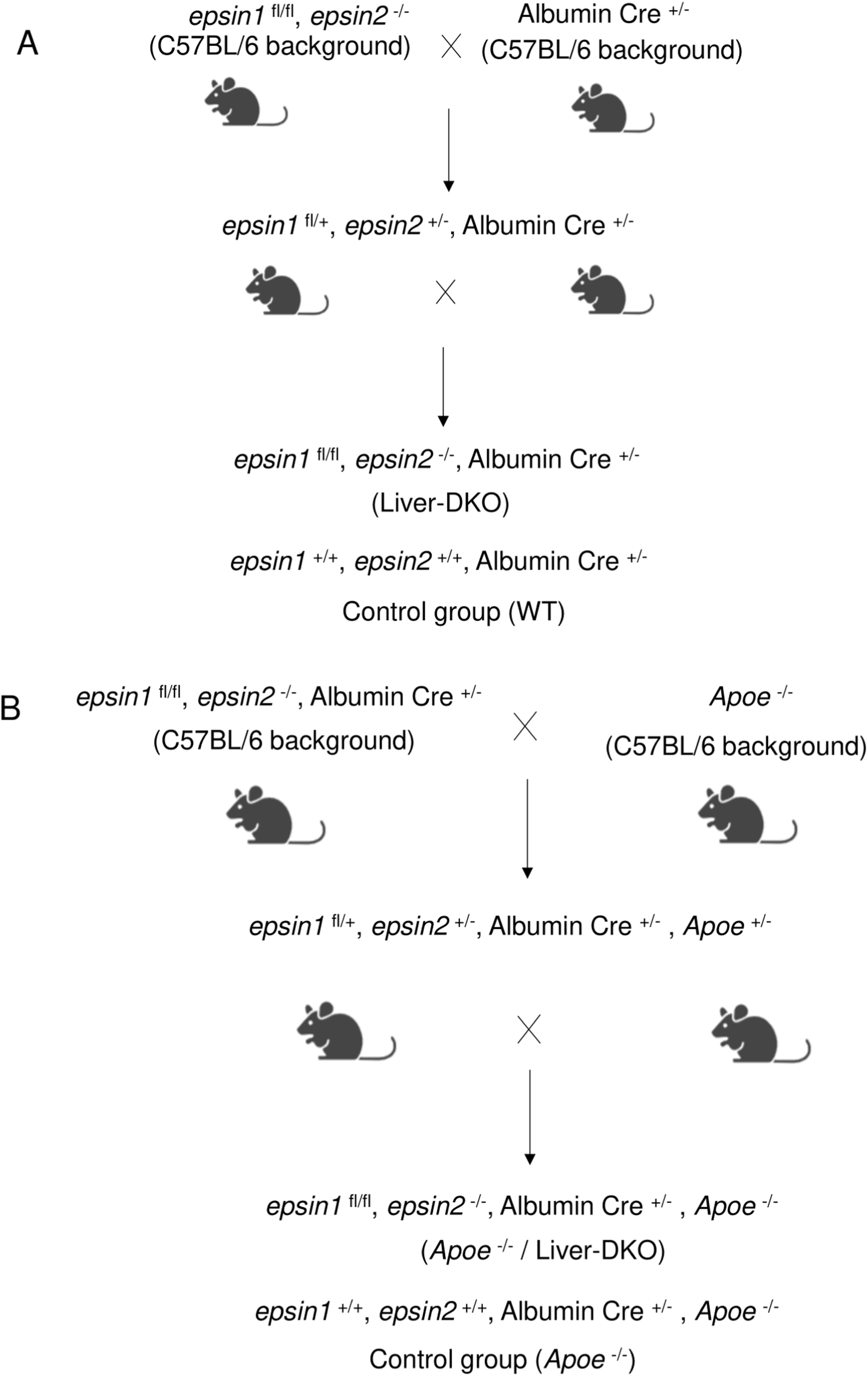
Overview of animal models. A: Workflow of generation of *epsin1* ^fl/fl^, *epsin2* ^-/-^, Albumin Cre ^+/-^ (Liver-DKO), *epsin1* ^+/+^, and *epsin2* ^+/+^, Albumin Cre ^+/-^ as control group (WT). B: Workflow of generation of *epsin1* ^fl/fl^, *epsin2* ^-/-^, Albumin Cre ^+/-^, *Apoe*^-/-^ (*Apoe* ^-/-^ /Liver-DKO), and *epsin1* ^+/+^, *epsin2* ^+/+^, Albumin Cre ^+/-^, *Apoe*^-/-^ as control group (*Apoe*^-/-^).

**Figure S2:**
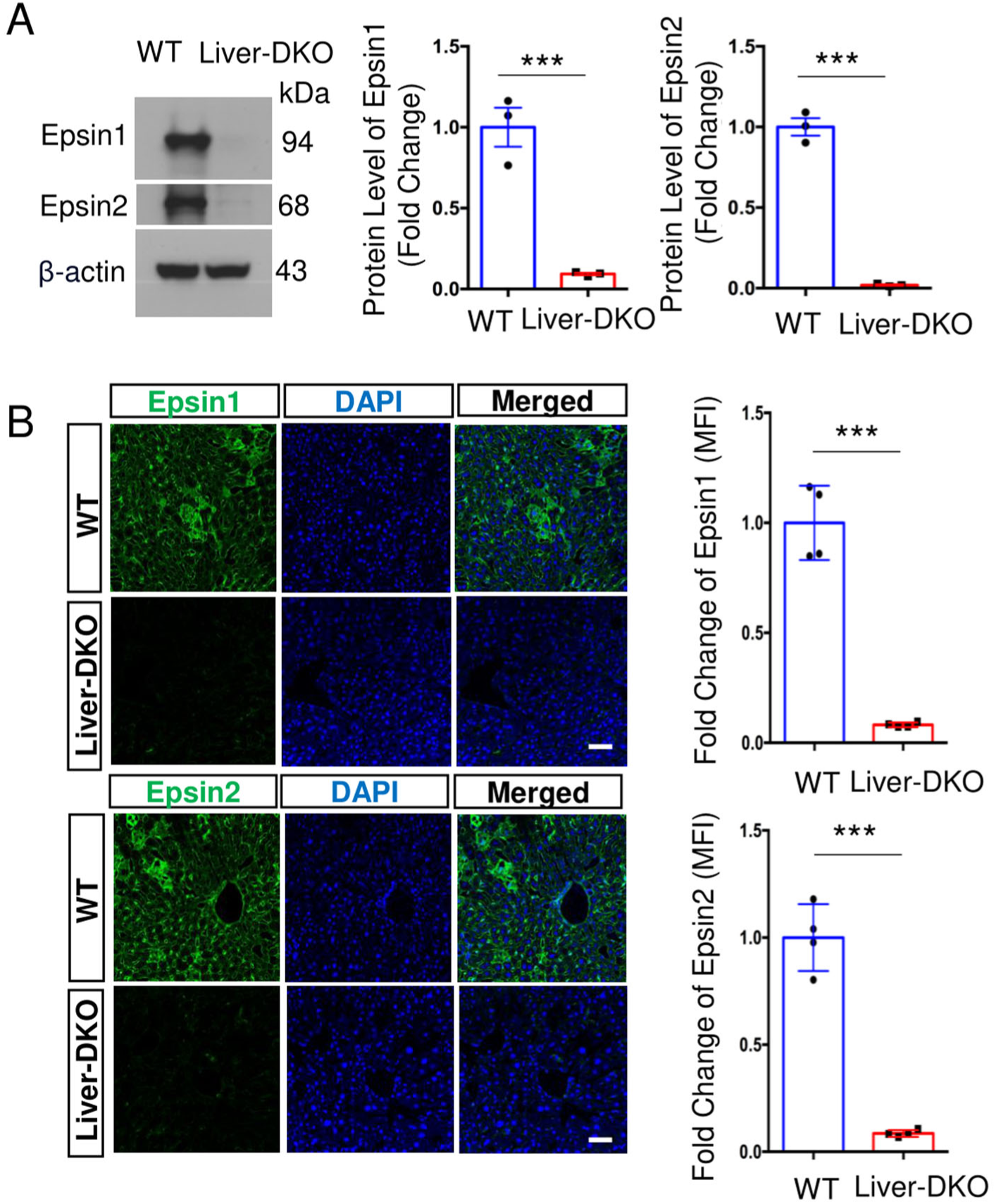
Evidence of the deficiency of Epsin1 and Epsin2 in Liver-DKO mice. A:Western blot shows no protein expression of both Epsin1 and Epsin2 proteins in the liver lysates isolated from Liver-DKO mice (left). Quantification of Epsin1 and Epsin2 protein level (right) (n=3, *** p<0.001). B: Immunofluorescence analyses validate no Epsin1 and Epsin2 protein expression in the liver cryosection from Liver-DKO mice (left). Quantification of the MFI of Epsin1 and Epsin2 (right) (n=5, *** p<0.001). Statistical analysis (A and B) in WT and Liver-DKO mice comparison is conducted by the Student’s t-test.

**Figure S3:**
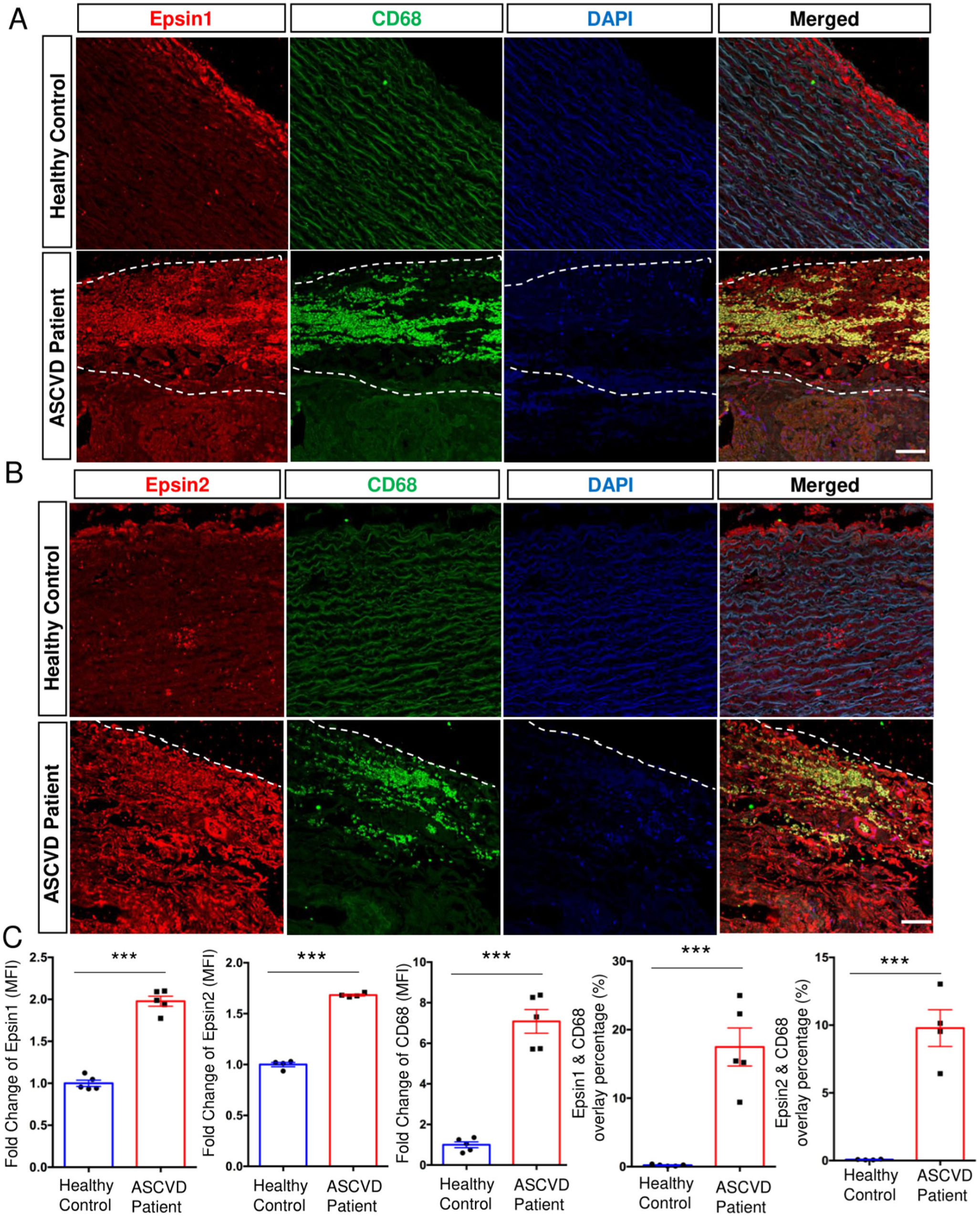
Elevated epsin1 and epsin2 expression in the aortic biopsies of ASCVD patients, and recruitment of CD68 positive macrophages in the aortas from ASCVD patients that colocalize with both epsin1 and epsin2. A: Immunofluorescence co-stain of Epsin1 and CD68 antibodies in the aortas of both healthy control and ASCVD patients. Epsin1 is red, CD68 is green, and DAPI is used for nuclear stains. The atherosclerotic lesion is encircled with a dashed line in ASCVD patients. Scale bar, 50 μm. B: Immunofluorescence co-stain of Epsin2 and CD68 antibodies in the aortas of both healthy control and ASCVD patients. Epsin2 is red, CD68 is green, and DAPI is used for nuclear stains. The atherosclerotic lesion is highlighted below the dashed line in ASCVD patients. Scale bar, 50 μm. C: Quantification of mean fluorescence intensity (MFI) of Epsin1, Epsin2 and CD68 proteins between healthy control and ASCVD patients. CD68 expression is highly colocalized with both Epsin1 and Epsin2 in ASCVD patients, and the overlay percentage between CD68 and Epsin1 or CD68 and Epsin2 are quantified (n=5, *** p<0.001). Statistical analysis (C) in healthy control and ASCVD patients’ comparison is conducted by the Student’s t-test.

**Figure S4:**
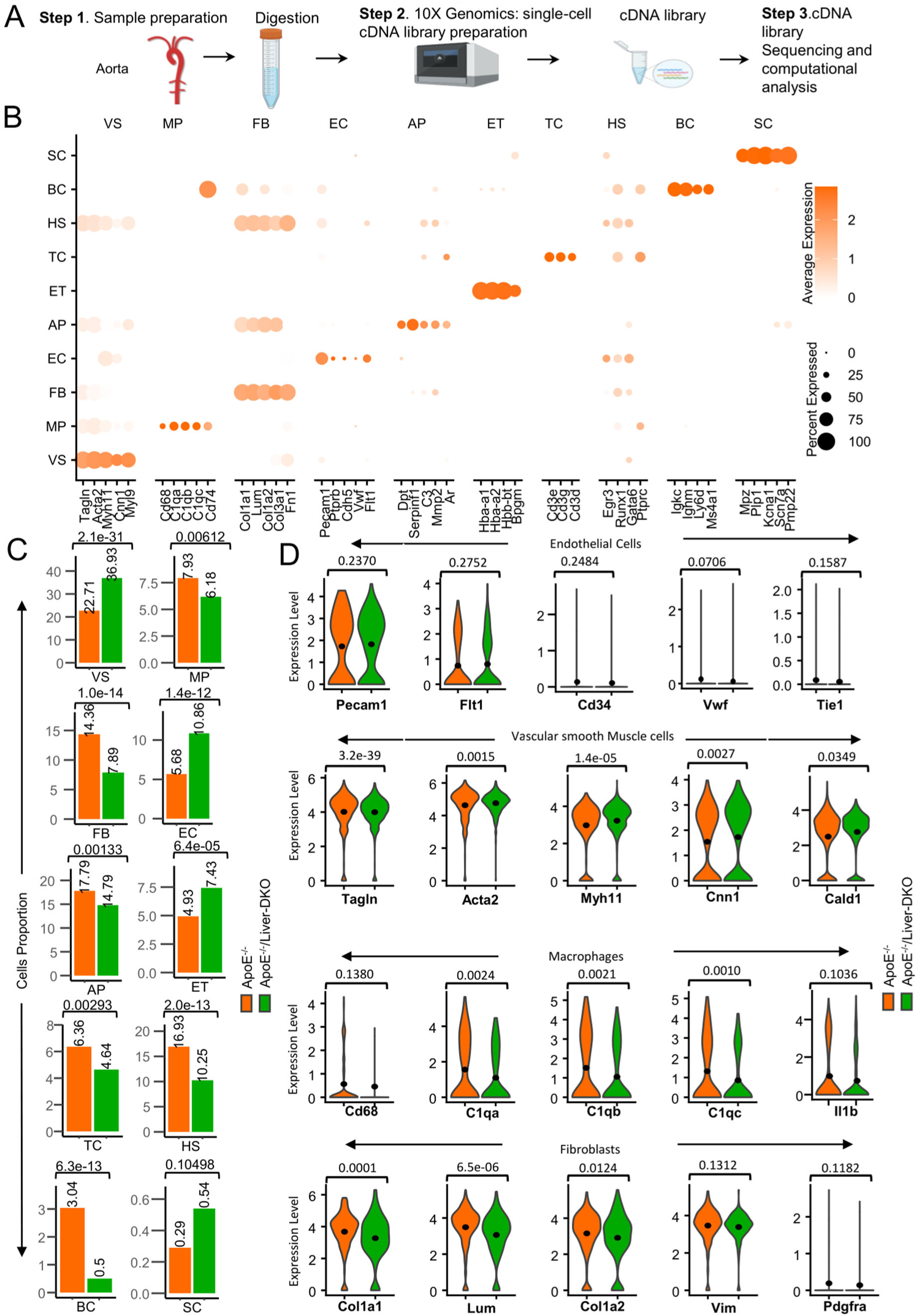
Single-cell RNA-sequencing reveals different cell types in the aortas in both ApoE^-/-^ controls and ApoE^-/-^/Liver-DKO mice. A: Schematic representation of the single-cell RNA-sequencing (scRNA-seq) process performed on aortas from ApoE^-/-^ and ApoE^-/-^/Liver-DKO mice. B: Dot plot illustrates the percentage of cells expressing each gene marker corresponding to specific cell types in ApoE^-/-^/Liver-DKO and ApoE^-/-^ mice. The size of the dots represents the proportion of cells expressing the marker, while the color intensity indicates the expression level of the gene marker in each cell type. C: Bar plots showing the cell proportions of different sub-cell types in the aortas of ApoE^-/-^ and ApoE^-/-^/Liver-DKO mice. A relatively higher proportion of VS, EC and a relatively lower proportion of MP, FB in the aortas of ApoE^-/-^/Liver-DKO than ApoE^-/-^ controls. D: Violin plots of markers of different cell types, including endothelial cells, vascular smooth muscle cells, macrophages, and fibroblasts. Endothelial cell markers: *Pecam1*, *Flt1*, *CD34*, *Vwf*, *Tei1*. Vascular smooth muscle cell makers: *Tagln*, *Acta2*, *Myh11*, *Cnn1*, and *Cald1*. Macrophage markers: *Cd68*, *C1qa*, *C1qb*, *C1qc*, *Il1b*. Fibroblast markers: *Col1a1*, *Lum*, *Col1a2*, *Vim*, *Pdgfra*. All p values of violin plots are labeled above. Statistical analysis (C, D) in ApoE^-/-^ and ApoE^-/-^/Liver-DKO comparison is conducted by the Student’s t-test.

**Figure S5:**
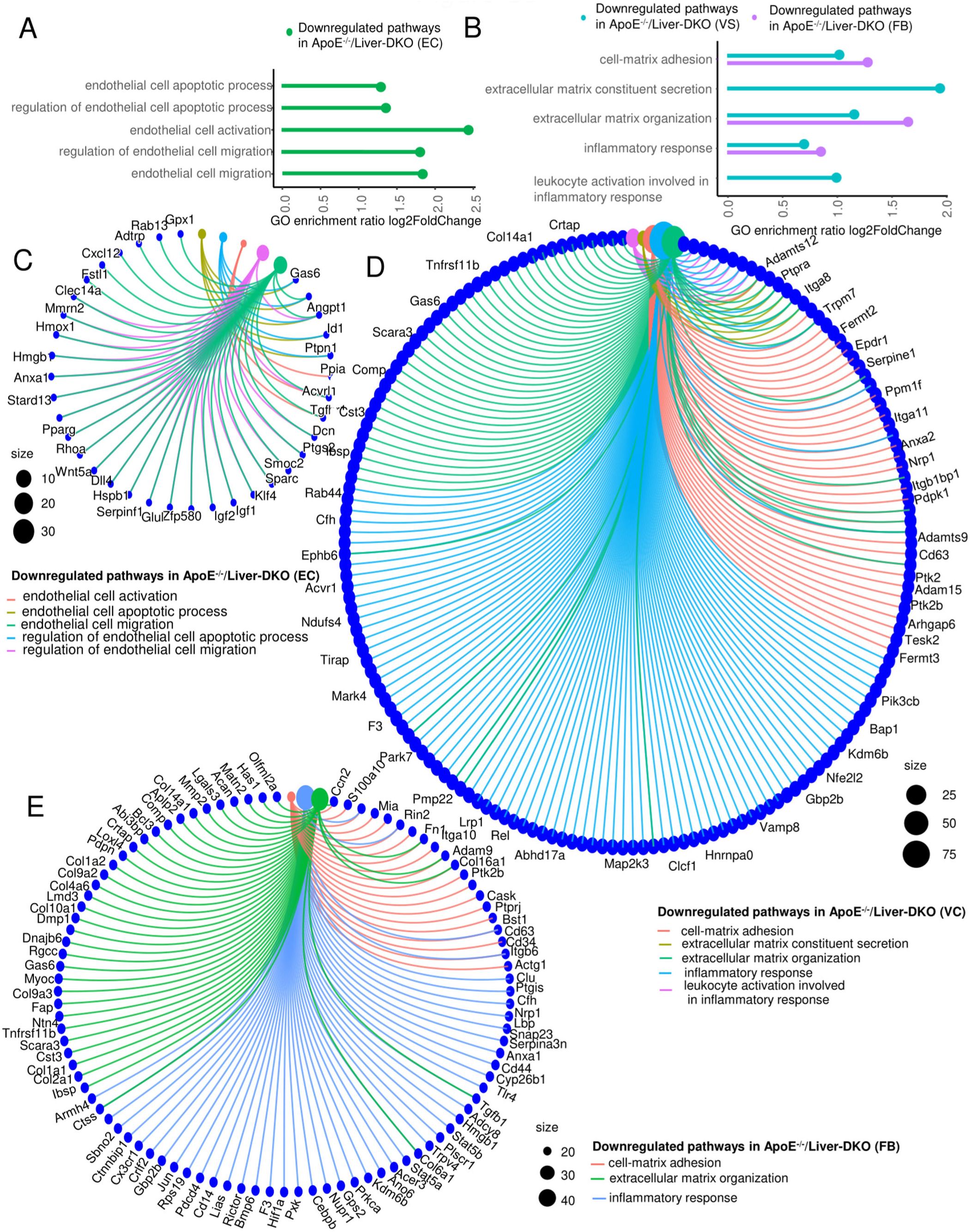
Single-cell RNA sequencing reveals upregulated or downregulated pathways in different cell types in the aortas in both ApoE^-/-^ controls and ApoE^-/-^/Liver-DKO mice. A-B: Gene Ontology (GO) analysis showing downregulated enriched pathways of regulation of endothelial cell apoptosis, endothelial cell activation, and endothelial cell migration in endothelial cells (A) and downregulated enriched pathways of regulation of cell-matrix adhesion, inflammatory response in both vascular smooth muscle cells and fibroblasts (B) in ApoE^-/-^/Liver-DKO mice relative to ApoE^-/-^ controls. C: Circular cnetplot illustrating significantly downregulated pathways and corresponding genes in endothelial cells (EC), such as endothelial cell activation, migration, and apoptotic process. D: Circular cnetplot representing downregulated pathways and corresponding genes in vascular smooth muscle cells (VS), including cell-matrix adhesion, extracellular matrix organization, and inflammatory response. E: Circular cnetplot representing downregulated pathways and corresponding genes in fibroblasts (FB), such as cell-matrix adhesion, extracellular matrix organization, and inflammatory response.

**Figure S6:**
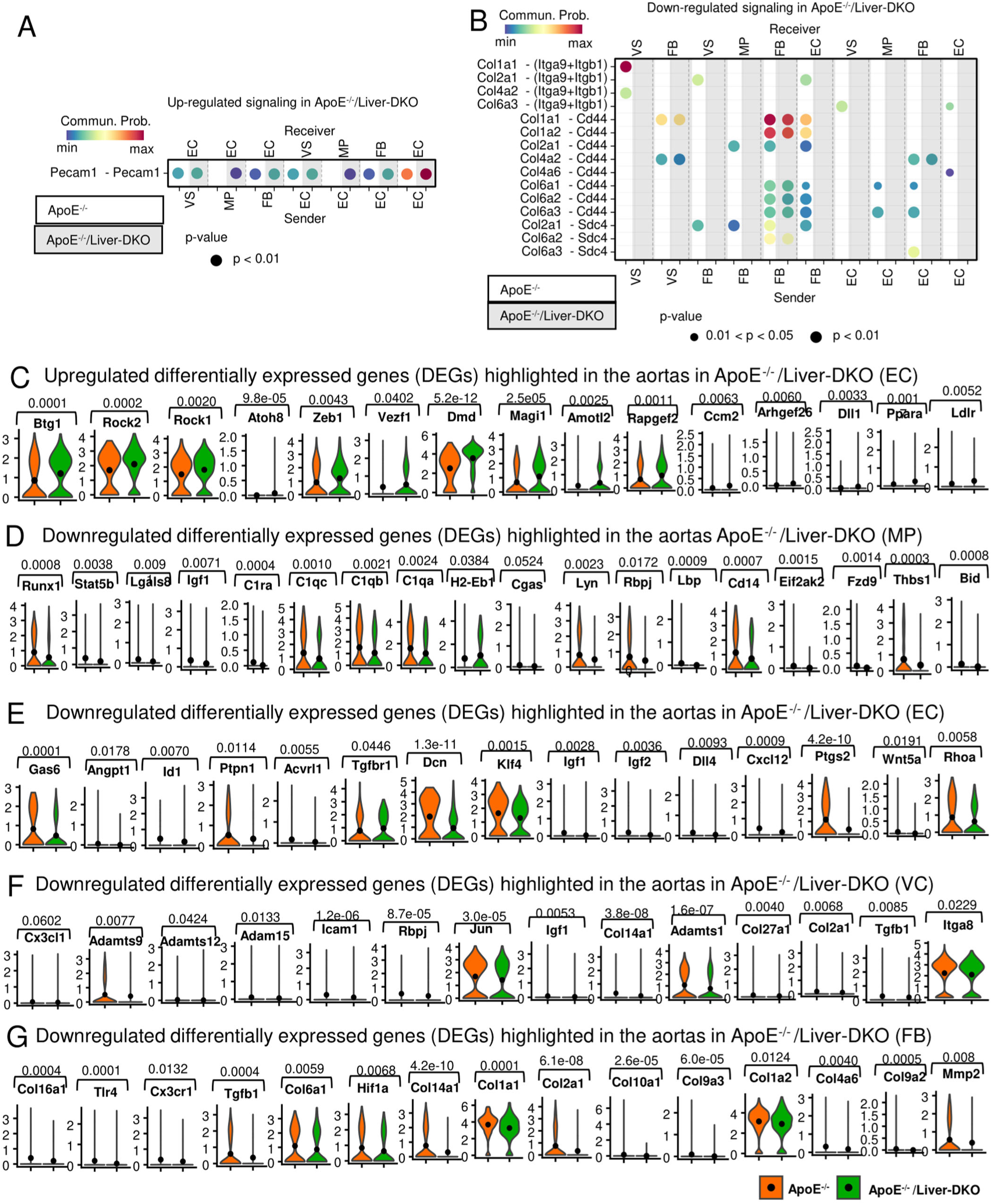
Single-cell RNA-sequencing upregulated and downregulated pathways in ApoE^-/-^/Liver-DKO, and violin plots highlighted the differentially expressed genes presented in cnet plots. A-B: Illustration of enhanced Pecam1-associated communication pathway (A) and inhibited Collagen-associated communication pathways (B) in ApoE^-/-^/Liver-DKO relative to ApoE^-/-^ controls, indicating diminished collagen synthesis in the aortas in ApoE^-/-^/Liver-DKO mice. C-G: Violin plots of differentially expressed genes (DEGs) that are highlighted in cnet plots of upregulated pathways in endothelial cells (EC) (C), downregulated pathways in macrophages (MP) (D), downregulated pathways in endothelial cells (EC) (E), downregulated pathways in vascular smooth muscle cells (VC) (F), and downregulated pathways in fibroblasts (FB) (G) in the aortas in ApoE^-/-^/Liver-DKO mice. Statistical analysis (C-G) in ApoE^-/-^ and ApoE^-/-^/Liver-DKO comparison is conducted by the Student’s t-test.

**Figure S7:**
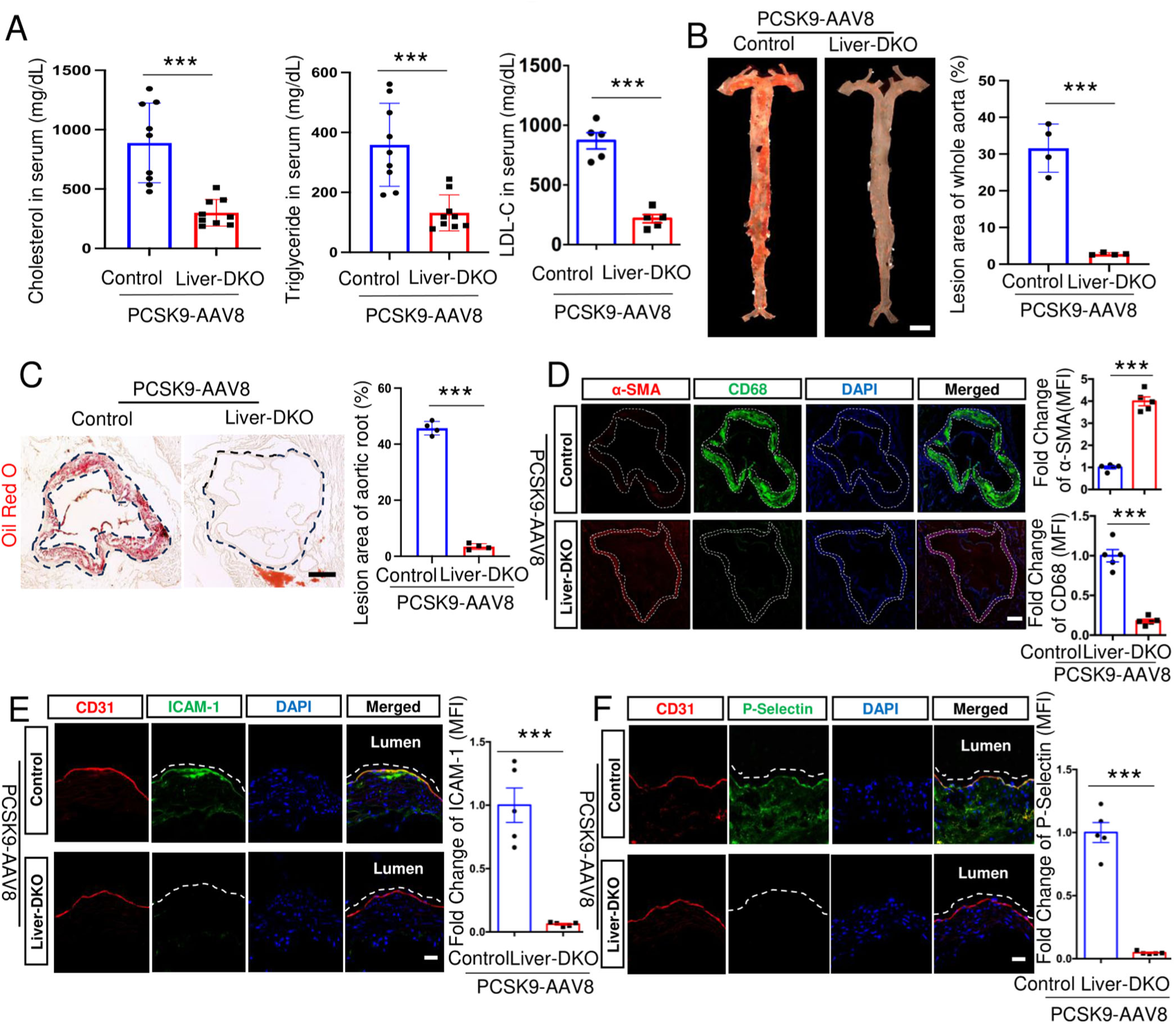
Depletion of epsins in the liver negates the effects of PCSK9 overexpression on atherosclerotic lesion formation, macrophage accumulation, and plasma lipid levels. A: Serum cholesterol, triglyceride, and low-density lipoprotein cholesterol (LDL-C) levels in WD-fed WT and Liver-DKO mice treated with PCSK9-AAV8 after 20 weeks on a WD (n=5-9, p***<0.001). B: *En face* ORO staining of aortas (left) from PCSK9-AAV8 injected WT or Liver-DKO mice fed a WD, and unpaired t-test (right) for the lesion areas (n=4, ***p< 0.001). Scale bar, 5 mm. C: Aortic roots from WD-fed PCSK9-AAV8 injected WT or Liver-DKO mice stained with ORO (left), and unpaired t-test (right) for the lesion areas (n=4, ***p< 0.001). Scale bar, 500 μm. D: Aortic roots from WD-fed PCSK9-AAV8 injected WT or Liver-DKO mice stained with CD68 macrophage marker and α**-**SMA vascular smooth muscle marker (left), and unpaired t-test (right) for quantification of the mean fluorescence intensity (MFI) (n=5, ***p< 0.001). Scale bar, 200 μm. E-F: IF staining of endothelial cell marker CD31 and endothelial cell adhesion molecules, such as ICAM-1 (E) and P-Selectin (F) (left), in the aortas between WD-fed PCSK9-AAV8 injected WT or Liver-DKO mice. The fold changes of ICAM-1 (MFI) and P-Selectin (MFI) were quantified (right) (n=5, ***p<0.001). Scale bar, 20 μm. Diminished expression of ICAM-1 and P-Selectin proteins in the aortas in WD-fed PCSK9-AAV8-induced Liver-DKO. All statistical analysis (A-F) in PCSK9-AAV8 injected WT and PCSK9-AAV8-induced Liver-DKO comparison is conducted by the Student’s t-test.

**Figure S8:**
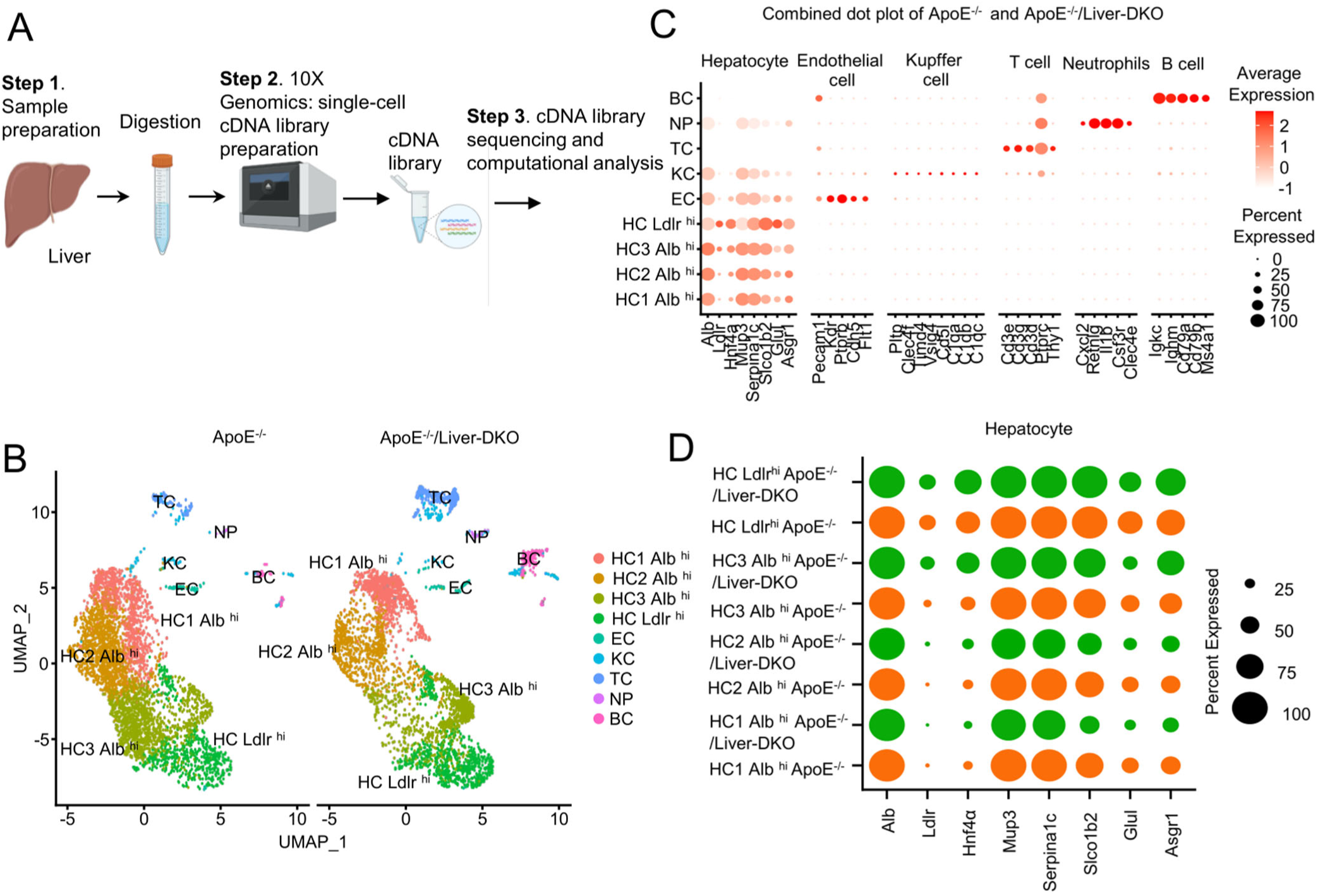
**Single-cell RNA-sequencing reveals different hepatocyte subtypes in the livers of ApoE^-^**^/-^ **and ApoE^-/-^/Liver-DKO mice.** A: Schematic representation of the single-cell RNA-sequencing (scRNA-seq) process performed on the livers of ApoE^-/-^ and ApoE^-/-^/Liver-DKO mice. B: UMAP visualization of whole cell populations in ApoE^-/-^/Liver-DKO and ApoE^-/-^ mice. C: Dot plot illustrates the percentage of cells expressing each gene marker corresponding to specific cell types in ApoE^-/-^/Liver-DKO and ApoE^-/-^ mice. The size of the dots represents the proportion of cells expressing the marker, while the color intensity indicates the expression level of the gene marker in each cell type. D: Dot plot of subclusters of hepatocytes according to the expression of hepatocyte markers, such as Albumin and Ldlr, in ApoE^-/-^/Liver-DKO and ApoE^-/-^ mice. HC1 Alb ^hi^, HC2 Alb^hi^, HC3 Alb^hi^, and HC Ldlr^hi^ hepatocytes are clustered.

**Figure S9:**
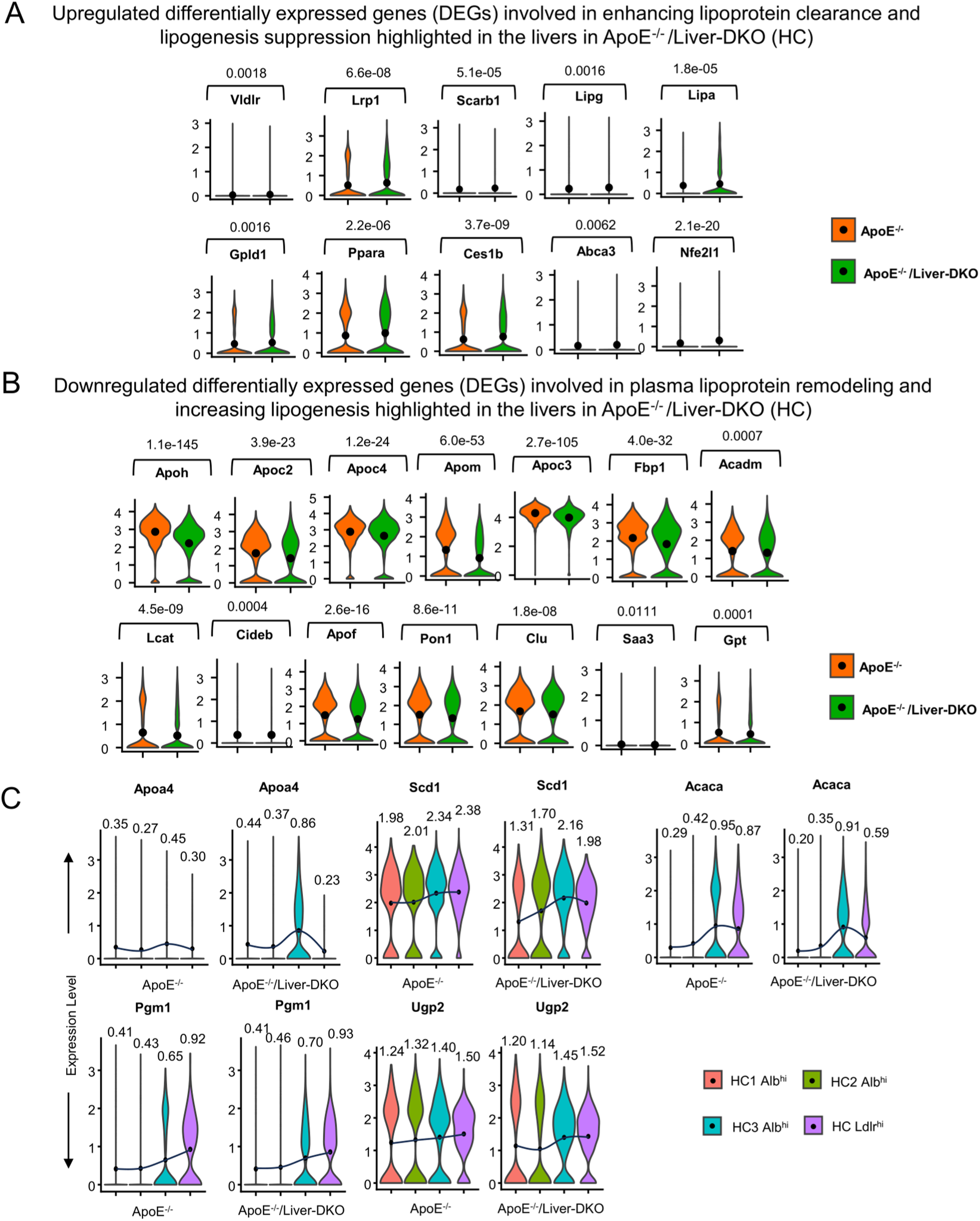
Elevated differentially expression genes (DEGs) that enhance low-density lipoprotein particle clearance, glycogenesis and lipogenesis suppression and diminished DEGs that facilitate hepatic lipogenesis and plasma lipoprotein particle assembly in the livers of ApoE^-/-^/Liver-DKO mice. A: Upregulated DEGs that increase low-density lipoprotein particle clearance and/or reduce lipogenesis in hepatocytes from ApoE^-/-^ /Liver-DKO mice. B: Downregulated DEGs that mediate plasma lipoprotein particle remodeling and hepatic lipogenesis are highlighted in hepatocytes from ApoE^-/-^ /Liver-DKO mice. C: Downregulated expression levels of lipogenic genes, such as *Scd1* and *Acaca*, and upregulated expression of *Apoa4* gene in the hepatocyte subpopulations from ApoE^-/-^/Liver-DKO mice, indicate diminished lipogenesis in the liver of ApoE^-/-^/Liver-DKO mice. Upregulated expression levels of glycogenic genes, such as *Pgm1* and *Ugp2*, in different clustered hepatocytes in the liver of ApoE^-/-^/Liver-DKO mice reveal elevated glycogenesis in the liver of ApoE^-/-^/Liver-DKO mice. The mean expression levels of genes are highlighted in the violin plots (p<0.05). All statistical analysis (A-C) in ApoE^-/-^ and ApoE^-/-^/Liver-DKO comparison is conducted by the Student’s t-test.

**Figure S10:**
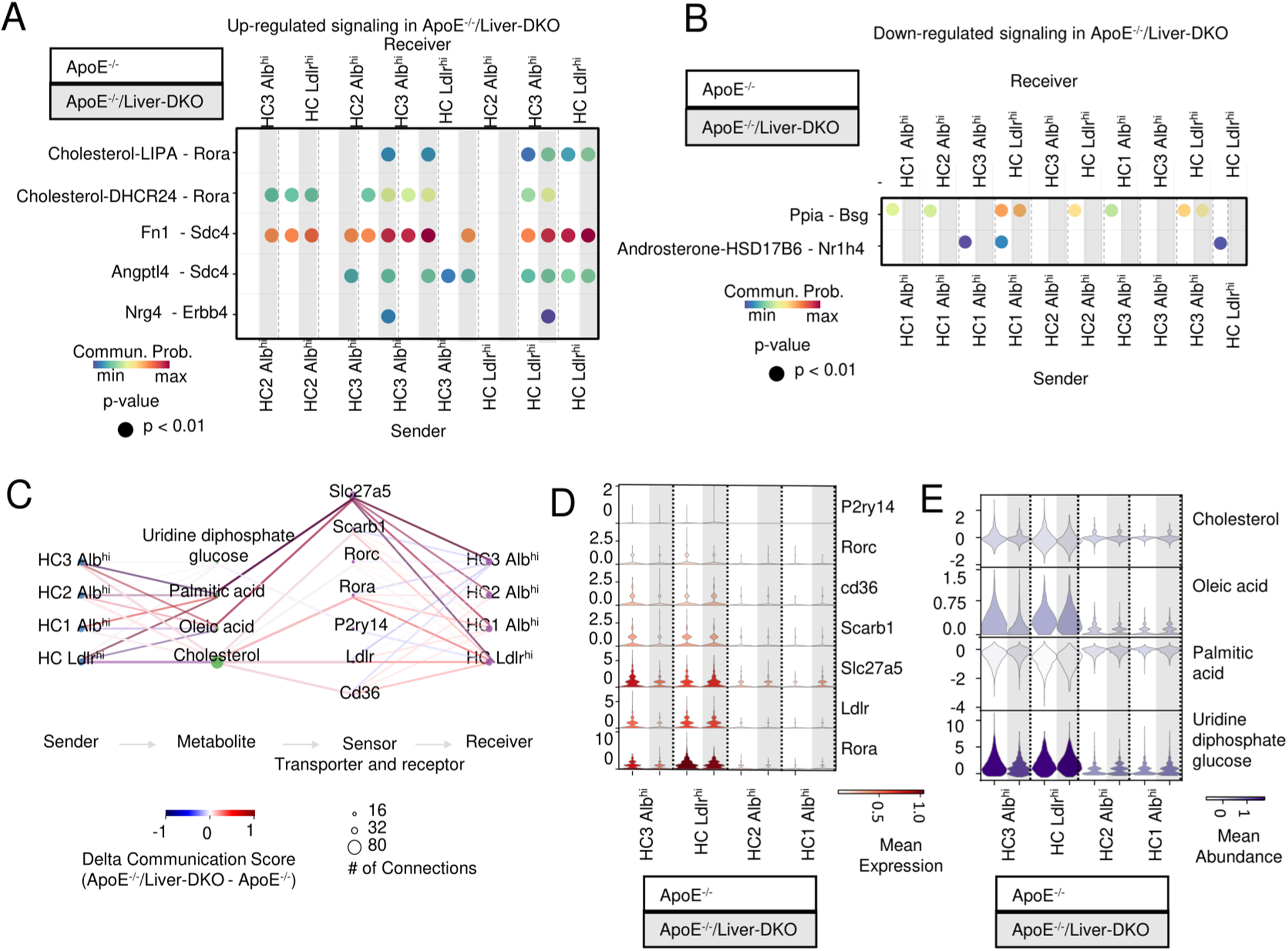
Enhanced LDL particle clearance and LDLR-cholesterol communication elucidate improved LDL-C clearance in ApoE^-/-^/Liver-DKO mice. A-B: Enhanced Rorα-cholesterol and Sdc4-Fn1 communication pathways (A) and diminished Bsg-Ppia and Nr1h4-AndrosteroneHSD17B6 communication pathways (B) in ApoE^-/-^/Liver-DKO hepatocytes. C: Quantitative analysis demonstrates metabolite communication score in ApoE^-/-^/Liver-DKO mice compared with ApoE^-/-^ controls, showing elevated communication score between cholesterol and Ldlr in ApoE^-/-^/Liver-DKO mice. D: Relative receptor expression in ApoE^-/-^/Liver-DKO mice compared with ApoE^-/-^ controls. E: Relative metabolite abundance in ApoE^-/-^/Liver-DKO mice compared with ApoE^-/-^ controls.

**Figure S11:**
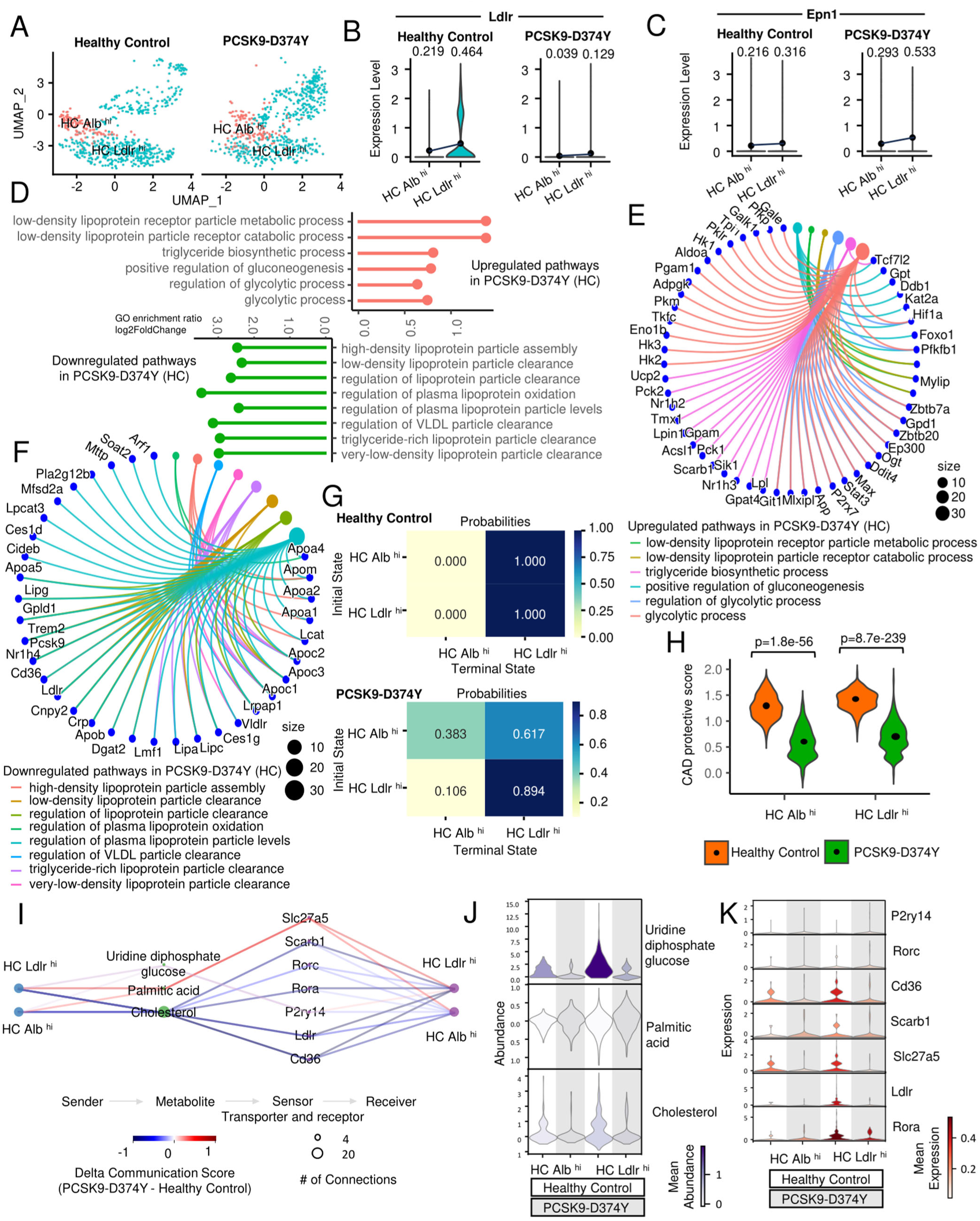
Single-cell RNA-sequencing reveals diminished expression of LDLR in the livers in PCSK9-D374Y mutated mice. A: UMAP visualization illustrating the heterogeneity of hepatocyte cell populations, showing distinct clustering patterns of hepatocytes in healthy control mice compared with PCSK9-D374Y mutants. B: Downregulated Ldlr expression in hepatocytes in PCSK9-D374Y mutant compared to healthy control. The mean expression levels of Ldlr are highlighted in the violin plots (p<0.05). C: Upregulated Epn1 expression in hepatocytes in PCSK9-D374Y mutant compared to healthy control. The mean expression levels of Epn1 are highlighted in the violin plots (p<0.05). D: Gene Ontology (GO) analysis showing significantly enriched pathways for upregulated glycolytic process and downregulated LDL particle clearance in PCSK9-D374Y mutants relative to healthy control. E: Circular cnetplot representing upregulated pathways and corresponding genes in PCSK9-D374Y hepatocytes (HC). F: Circular cnetplot illustrating significantly downregulated pathways and corresponding genes in PCSK9-D374Y hepatocytes (HC). G: CellRank analysis illustrating inhibited propensity from Alb^hi^ hepatocytes to Ldlr^hi^ hepatocytes in PCSK9-D374Y mutants compared to healthy controls. H: Coronary artery disease (CAD) protective score comparing PCSK9-D374Y mutant with healthy control, showing a lower CAD protective score in PCSK9-D374Y mutant than healthy control. I: Quantitative analysis demonstrating metabolite communication scores in PCSK9-D374Y-mutated mice compared with healthy controls, showing diminished communication scores between cholesterol and Ldlr in PCSK9-D374Y mutants. J: Relative metabolite expression abundance in PCSK9-D374Y mutant compared to healthy control, showing downregulated expression of uridine diphosphate glucose (UDPG) in PCSK9-D374Y mutant compared to healthy control. K: Relative metabolite receptor expression abundance in PCSK9-D374Y mutant compared with healthy control, revealing diminished expression of Ldlr in PCSK9-D374Y mutant compared to healthy control. Statistical analysis (B, C, H) in Healthy Control and PCSK9-D374Y comparison is conducted by the Student’s t-test.

**Figure S12:**
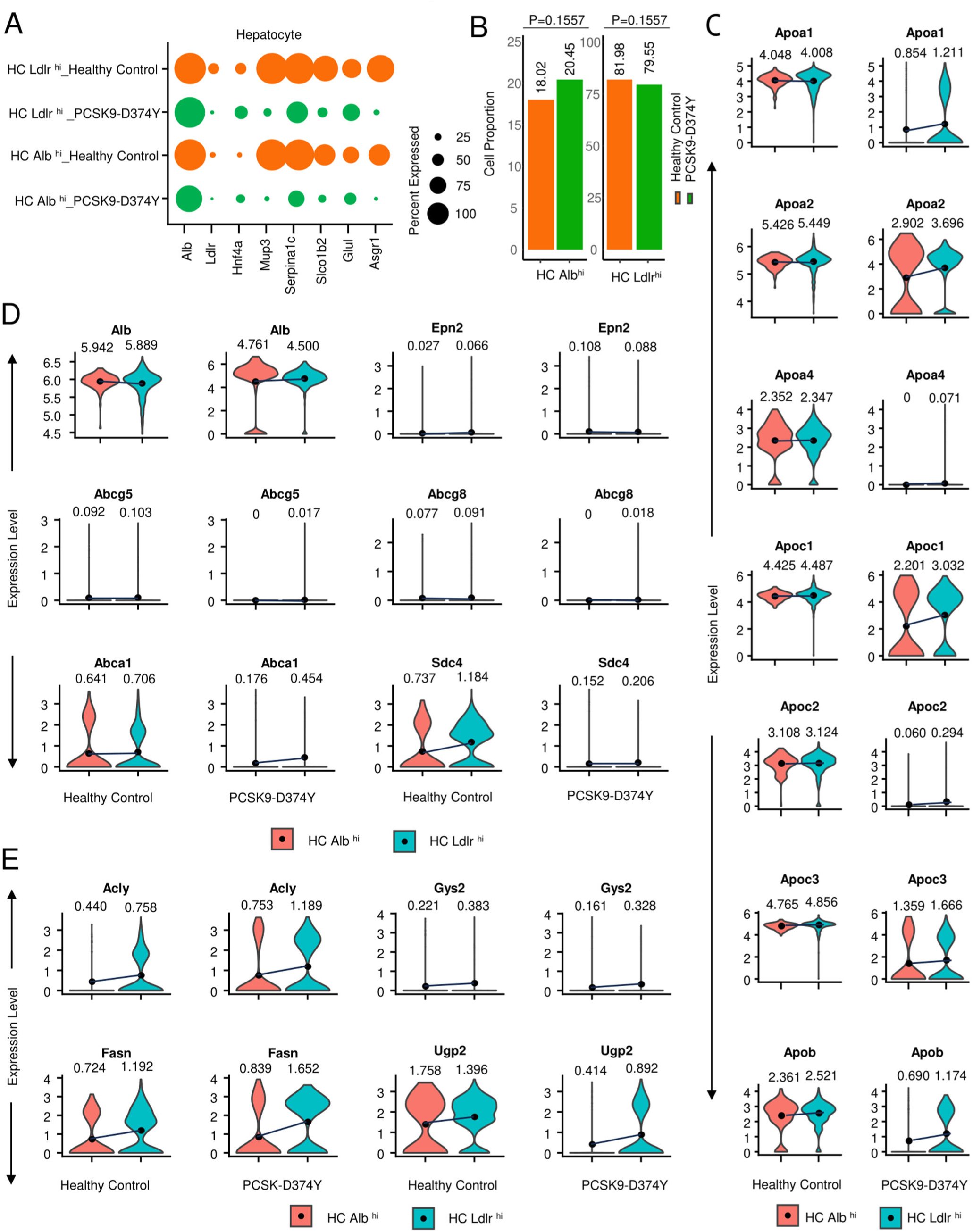
Reduced Ldlr^hi^ hepatocytes in the liver, enhanced gene expression of lipogenesis and diminished expression of genes involved in glycogenesis in PCSK9-D374Y mutant compared with healthy control. A: Dot plot of subclusters of hepatocytes according to the expression of hepatocyte markers, such as Albumin and Ldlr, in healthy control and PCSK9-D374Y mice. B: A relatively low proportion of Ldlr^hi^ hepatocytes in the livers of PCSK9-D374Y mice compared with healthy control. C: Downregulated expression of apolipoprotein genes, such as *Apoa1*, *Apoa2*, *Apoa4*, *Apoc1*, *Apoc2*, *Apoc3* and *Apob*, in different clustered hepatocytes (HC Alb^hi^, HC Ldlr^hi^) in PCSK9-D374Y mutant compared with healthy control, indicates diminished low-density lipoprotein clearance in PCSK9-D374Y mutant. The mean expression levels of apolipoprotein genes are highlighted in the violin plots (p<0.05). D: Downregulated expression levels of genes involved in cholesterol efflux, such as *Abca1*, *Abcg5*, *Abcg8*, and genes that participate in fatty acid metabolism (*Sdc4*) in the different clustered hepatocytes in the liver from PCSK9-D374Y mutant, indicates the dyslipidemia in PCSK9-D374Y mutant. Diminished expression of *Alb* gene in the different clustered hepatocytes (HC Alb^hi^, HC Ldlr^hi^) in the liver in PCSK9-D374Y mutant. Elevated expression of *Epn2* in both Alb^hi^ and Ldlr^hi^ hepatocytes in the liver in PCSK9-D374Y mutant. The mean expression levels of genes are highlighted in the violin plots (p<0.05). E: Upregulated expression of genes involved in lipogenesis, such as *Acly* and *Fasn*, in the different clustered hepatocytes in the liver in PCSK9-D374Y mutant. Downregulated expression of genes that participate in glycogenesis, such as *Gys2* and *Ugp2*, in the different clustered hepatocytes in the liver in PCSK9-D374Y mutant. The mean expression levels of genes are highlighted in the violin plots (p<0.05). Statistical analysis (B, C, D, E) in Healthy Control and PCSK9-D374Y comparison is conducted by the Student’s t-test.

**Figure S13:**
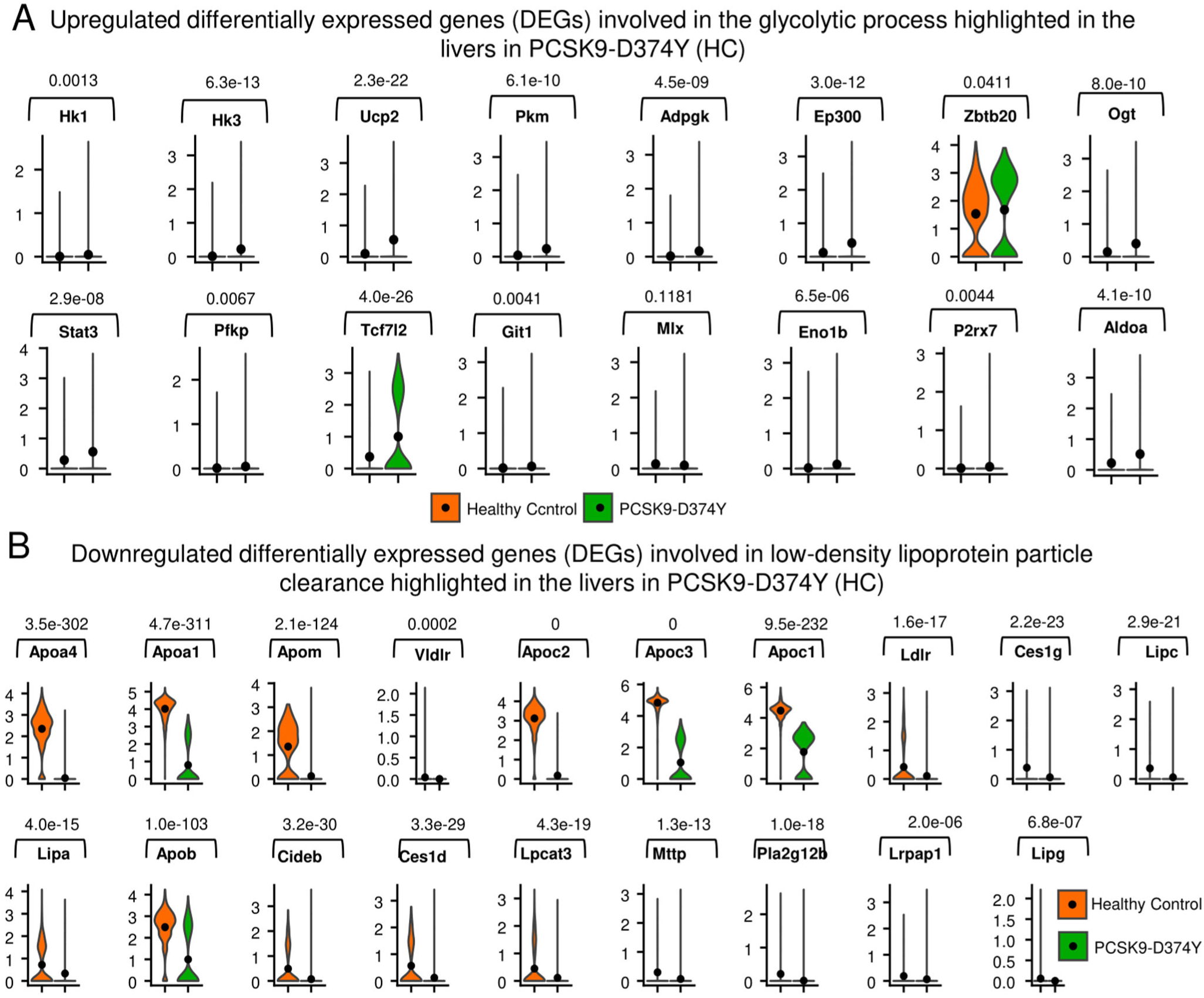
Elevated expression of genes involved in the glycolytic process, and diminished expression of genes responding to low-density lipoprotein particle clearance in PCSK9-D374Y mutant. A-B: Violin plots of differentially expressed genes (DEGs) that are highlighted in cnet plots of either upregulated glycolytic process pathways (A) or downregulated low-density lipoprotein particle clearance pathways (B) in the hepatocytes (HC) in PCSK9-D374Y mutant (p<0.05). All statistical analysis (A and B) in Healthy Control and PCSK9-D374Y comparison is conducted by the Student’s t-test.

**Figure S14:**
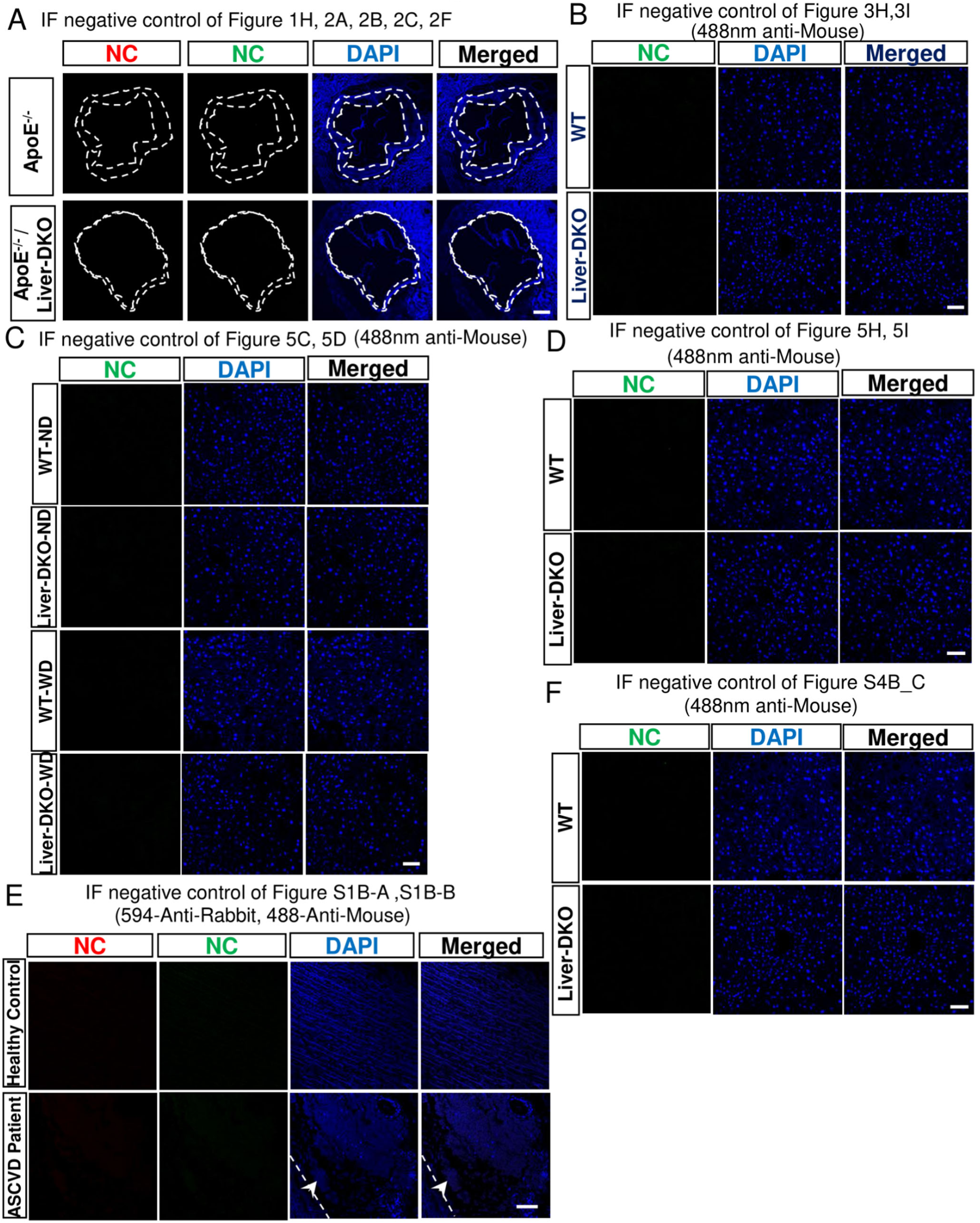
Negative controls of immunofluorescence (IF) images without primary antibody staining. A-F: the negative controls of mouse aortas (A) (scale bar, 200 μm), mouse livers (B, C, D, F) (scale bar, 50 μm), and human aortic samples (E) (scale bar, 50 μm), which are annotated as negative controls of either main IF figures or supplemental IF figures, respectively.

**Table S1.** Genes harboring variants for coronary artery disease (CAD) in ClinVar.

| Phenotypes or Disorders | Genes |
| --- | --- |
| Abnormality of lipid metabolism | ApoE |
| Hypocholesterolemia | ApoB, LDLR |
| Hypobetalipoproteinemia | ApoB |
| Familial hypoalphalipoproteinemia | ApoA1 |
| Hyperalphalipoproteinemia | ApoC1 |
| Hyperapobetalipoproteinemia | LPL |
| Hyperlipoproteinemia | LPL |
| Familial hypertriglyceridemia | ApoA5 |
| Lysosomal acid lipase deficiency | LIPA |
| High density lipoprotein cholesterol level<br>quantitative trait locus | SCARB1 |
| Cholesteryl ester storage disease | LIPA |
| Wolman disease | LIPA |
| Sitosterolemia | ABCG5, ABCG8 |

**Table S2.** Human aorta samples for both coronary artery disease patients and healthy controls.

| Individual name | Gender | Tissue Source | Normal/ Diseased | Age | Specific Diagnosis | Disease Type |
| --- | --- | --- | --- | --- | --- | --- |
| RA06-1040 | M | Aorta | Diseased | 62-83 | Atherosclerosis | Atherosclerotic lesion |
| RA06-2816 | M | Aorta | Diseased | 62-83 | Atherosclerosis | Atherosclerotic lesion |
| RA03-0422 | F | Aorta | Diseased | 62-83 | Atherosclerosis | Atherosclerotic plaque with calcifications |
| RA02-0678 | M | Aorta | Diseased | 62-83 | Atherosclerosis | Mild atherosclerotic change |
| RA03-0488 | M | Aorta | Diseased | 62-83 | Atherosclerosis | Thrombus and atherosclerotic plaque severe |
| RA03-0857 | F | Aorta | Diseased | 62-83 | Atherosclerosis | Atherosclerosis with subintimal hemorrhage |
| RA03-1686 | M | Aorta | Normal | <20 | Dissection | N/A |
| R14-0199 | M | Aorta | Normal | 62-83 | Dissection | N/A |
| RA04-2408 | F | Aorta | Normal | 62-83 | Degeneration | N/A |

